# IL-1β Signalling Modulates T Follicular Helper and Regulatory Cells in Human Lymphoid Tissues

**DOI:** 10.1101/2024.10.02.616222

**Authors:** Romain Vaineau, Raphaël Jeger-Madiot, Samir Ali-Moussa, Laura Prudhomme, Hippolyte Debarnot, Nicolas Coatnoan, Johanna Dubois, Marie Binvignat, Hélène Vantomme, Bruno Gouritin, Gwladys Fourcade, Paul Engeroff, Aude Belbézier, Romain Luscan, Françoise Denoyelle, Roberta Lorenzon, Claire Ribet, Michelle Rosenzwajg, Bertrand Bellier, David Klatzmann, Nicolas Tchitchek, Stéphanie Graff-Dubois

## Abstract

**Background:** Dysregulation of the T follicular helper (Tfh) and T follicular regulatory (Tfr) homeostasis in the germinal center (GC) can result in antibody-mediated autoimmunity. While interleukin-1β (IL-1β) has been shown to be an important modulator of the GC response in animal models via the expression of IL-1 agonist (IL-1R1) and antagonist (IL-1R2) receptors on follicular T cells, such regulation has not yet been studied in humans.

**Methods:** We investigated Tfh and Tfr phenotypes in human secondary lymphoid organs — namely tonsils, spleens, and mesenteric lymph nodes — using flow cytometry, single-cell transcriptomics, and *in vitro* cell culture. We also benchmarked our findings with a cohort of patients with autoimmune and inflammatory diseases.

**Results:** We found that Tfh and Tfr cells exhibit organ-specific phenotypes related to their activation status and IL-1 receptor expression. An excess of IL-1R1 over IL-1R2 was linked to the emergence of a unique activated Tfr subset that combines features of both Treg and GC-Tfh cells. Single-cell transcriptomics and *in vitro* studies showed that IL-1β signaling through IL-1R1 promotes follicular T-cell activation. Inhibiting IL-1β resulted in upregulation of IL-1R1 expression, showing a fine-tuned regulation. In autoimmune patients, high IL-1β and circulating Tfr levels correlated with higher autoantibody levels, linking inflammation, IL-1β signaling, and the Tfr/Tfh balance.

**Conclusions:** Our study underscores the pivotal role of IL-1β in follicular T-cell activation, contributing to pathological antibody production in humans. Targeting IL-1β signaling in Tfh and Tfr cells could offer new treatment strategies for antibody-mediated autoimmune diseases.

## Introduction

Germinal centers (GCs) are specialized structures enabling the production of high-affinity antibodies by B cells within secondary lymphoid organs (SLO)^1–3^. The GC reaction occurs following antigen-specific recognition, leading to the expansion of follicular helper T cells (Tfh) that support the maturation of cognate B cells^4–6^. Excessive help provided during the GC reaction can lead to pathologic antibody production, notably in the context of autoimmune diseases^7^. To maintain immune homeostasis, distinct regulatory mechanisms exist, encompassing external regulation via antibody feedback ^8^ or internal regulation through follicular regulatory T cells (Tfr)^9^.

Tfr are a suppressive population derived from thymic Treg^10^, which exert their functions by direct co-inhibition or by the secretion of immunomodulatory cytokines^11–13^. Accordingly, decreased circulating Tfr (cTfr) in the blood of patients are associated with disease onset and outcome in rheumatoid arthritis, systemic lupus erythematosus and multiple sclerosis^14–16^. However, emerging research questions the dichotomic view of Tfh and Tfr, by highlighting Tfr heterogeneity^17–19^ or by pointing out Tfh as precursors of Tfr^20–22^. Cytokines have a central role in the function of Tfh and Tfr, either by polarizing their differentiation^7^, or by guiding their migration such as CXCL13^23^, or else by serving as effector cytokines — i.e. IL-21 and IL-10 for Tfh and Tfr respectively^24,25^.

Interleukin-1β (IL-1β) is a proinflammatory cytokine with established roles in immune regulation, spanning both innate and adaptive responses^26^. Indeed, IL-1β has been shown to enable T-dependent antibody responses^27^, and to directly activate CD4^+^ T cells^28^. Notably, the expression of IL-1 receptors (IL-1Rs) including IL-1R1, the agonist, and IL-1R2, the decoy, have been associated with follicular T cell function. While the significance of IL-1β in various inflammatory processes is acknowledged, its interaction with follicular T cells (Tfol) and subsequent GC reaction is not fully understood. In mice, both binding to and inhibition of IL-1R1 impact Tfh activation and functions^29–32^. In particular, we previously demonstrated that the expression of IL-1R2 on murine Tfr cells inhibits IL-1R1-mediated activation of both Tfh and Tfr cells^32^. In humans, it has been documented that circulating Tfh (cTfh) are among the most responsive to IL-1β via NF-κB signaling^33^.

Given the difficult access to human lymphoid organs, most studies on the reactivity of Tfh and Tfr cells to IL-1β have been confined to murine studies or to circulating human Tfh and Tfr. In the present study, we aimed to understand the contribution of IL-1β to the activation profile of healthy human lymphoid-resident Tfh and Tfr. We also seek to decipher the role of IL-1β in the context of pathological antibody production.

## Materials and methods

### Collection of human secondary lymphoid organ samples

Tonsils were collected from immunocompetent children who underwent a partial tonsillectomy for the treatment of obstructive sleep apnea syndrome, at Necker-Enfants Malades Hospital, Paris, France. The collection and use of these samples were authorized by the ethics committee of Sorbonne University (CER-2021-043). Paired mesenteric lymph nodes and spleens were obtained from adult organ donors. Samples were obtained from the digestive surgery department at Pitié-Salpêtrière Hospital, Paris, France. The collection and use of these samples were authorized under protocol PFS-14-009 by the French Biomedicine Agency.

### Dissociation of secondary lymphoid organs and lymphoid cell isolation

Mesenteric lymph nodes, spleens, and tonsils were all treated following the same cell dissociation protocol. First, freshly collected samples were briefly washed with ethanol, sliced into pieces, and then transferred into C tubes (Miltenyi) containing complete medium: RPMI-1640 medium (Sigma-Aldrich) supplemented with 10% Fetal Bovine Serum (FBS), antibiotics (100 U Penicillin and 100 µg Streptomycin), and L-Glutamine. Cell dissociation was performed using a gentleMACS dissociator (Miltenyi), and the tube contents were filtered through 70 µm cell strainers. Mononuclear cells were isolated using a density gradient (Histopaque^®^-1077, Merck). Cell viability was assessed using trypan blue staining. The cell suspensions were frozen in cryotubes with FBS and 10% DMSO and stored at -80°C for short-term experiments and in liquid nitrogen for long-term preservation.

### Phenotypic profiling of human secondary lymphoid organs

Cell suspensions from mesenteric lymph nodes, spleens, and tonsils were thawed and left to rest overnight in a complete medium. The cells were then stained for viability using 405/420 Viobility (1/500, Miltenyi), washed and then stained with the following anti-human antibodies: CD3 (PerCP-Vio700, 1/100, Miltenyi), ICOS (Vioblue, 1/100, Miltenyi), CXCR5 (PE-Vio770, 1/500, Miltenyi), CD4 (APC-Vio770, 1/500, Miltenyi), CD38 (BV711, 1/400, BioLegend), CD25 (APC, 1/160, BioLegend), HLA-DR (BV786, 1/20, BD Biosciences), CD19 (PE-CF594, 1/200, BD Biosciences), IL-1R1 (PE, 1/80, R&D Systems), IL-1R2 (AF700, 1/80, R&D Systems), PD-1 (PE-Cy5.5, 1/320, Beckman Coulter). Staining was performed in 1X PBS with 5% FBS and incubated in the dark for 30 minutes. The cells were fixed using the BD Pharmingen Transcription Factor kit according to the manufacturer’s protocol. Intracellular staining was performed with the following anti-human antibodies: Ki67 (BUV395, 1/160, BD Biosciences), Foxp3 (AF488, 1/20, BD Biosciences), CTLA4 (BV605, 1/40, BioLegend). The cells were incubated for 40 minutes at 4°C, washed with the washing buffer, resuspended in 1X PBS, and analyzed using a CytoFLEX LX cytometer (Beckman Coulter).

### Functional profiling of human secondary lymphoid organs

Cell suspensions from mesenteric spleens and tonsils were thawed and left to rest overnight in a complete medium at 37°C and 5% CO_2_. The cells were stained with the following antibodies: CXCR5 (PE-Vio770, 1/500, Miltenyi) and CD4 (APC-Vio770, 1/500, Miltenyi). The cells were then fixed using the BD Pharmingen Transcription Factor kit. Intracellular staining was performed with the following anti-human antibodies: Ki67 (BUV395, 1/160, BD Biosciences), Foxp3 (AF488, 1/20, BD Biosciences), CD38 (BV711, 1/400, BioLegend), CD19 (PE-CF594, 1/200, BD Biosciences), IL-1R1 (PE, 1/80, R&D Systems), IL-1R2 (AF700, 1/80, R&D Systems), PD1 (PE-Cy5.5, 1/320, Beckman Coulter), IL-4 (PerCP-Cy5.5, 1/160, BD Biosciences), IL-21 (APC, 1/200, Miltneyi), IL-10 (BV421, 1/160, Biolegend), IL-2 (BV605, 1/160, Biolegend). The cells were incubated for 40 minutes at 4°C, washed with the washing buffer, resuspended in 1X PBS, and analyzed using a CytoFLEX LX cytometer (Beckman Coulter).

### *In vitro* activation of tonsillar whole cell suspensions

Frozen cell suspensions were thawed in complete medium, and washed twice, and live cells were counted using trypan blue staining. The cells were then incubated at 37°C and 5% CO_2_ overnight prior to cell activation. Whole-cell suspensions were centrifuged and resuspended in a complete medium with anti-CD3/CD28 activation beads (Dynabeads^TM^ Human T-Activator CD3/CD28, Thermo Fisher Scientific) at a concentration of 1 µL per 80,000 cells. Suspensions from each sample were then distributed into a 48-well plate with 500,000 cells per well. Three tonsillar conditions for cell activation were established: (1) stimulation alone, (2) stimulation with IL-1β premium grade (0.5 µg/mL, Miltenyi), and (3) stimulation with anti-IL-1 (Anakinra) (1 µg/mL, Kineret, SOBI). The plates were incubated at 37°C and 5% CO2 for 24 hours. After the incubation, cells were resuspended in 1X PBS prior to flow cytometry staining.

### Phenotypic profiling of cultured tonsillar whole cell suspensions

Both unstimulated cells (as controls) and cultured whole-cell suspensions were then stained for viability using 405/420 Viobility (1/500, Miltenyi), washed and then stained with the following anti-human antibodies: ICOS (Vioblue, 1/100, Miltenyi), CD3 (PerCP-Vio700, 1/100, Miltenyi), CXCR5 (PE-Vio770, 1/500, Miltenyi), CD19 (APC, 1/500, Miltenyi), CD4 (APC-Vio770, 1/500, Miltenyi), HLA-DR (BV786, 1/20, BD Biosciences), IL-1R1 (PE, 1/80, R&D Systems), IL-1R2 (AF700, 1/80, R&D Systems), CTLA-4 (BV605, 1/40, BioLegend), PD-1 (PE-Cy5.5, 1/320, Beckman Coulter). Staining was performed in 1X PBS with 5% FBS and incubated in the dark for 30 minutes. The cells were then washed and fixed using the BD Pharmingen Transcription Factor kit, and incubated for 45 minutes. Subsequently, the cells were washed with the kit washing buffer and stained with the intracellular mix containing the following anti-human antibodies: Ki67 (BUV395, 1/160, BD Biosciences) and Foxp3 (AF488, 1/20, BD Biosciences). The cells were incubated for 40 minutes at 4°C, washed again with the washing buffer, resuspended in 1X PBS, and analyzed using a Cytoflex LX cytometer (Beckman Coulter).

### Supervised analysis of flow cytometry data

Raw cytometry data were compensated using CytExpert software (Beckman Coulter). Manual gating was performed using FlowJo software (BD), from which geometric mean fluorescence intensity (MFI) and percentages relative to parent populations were extracted for each population of interest. Additional gates were created to determine marker positivity and extracted for each relevant cell population.

### Unsupervised analysis of flow cytometry data

Unsupervised analyses were performed on different cell populations, including CD4^+^ CXCR5^+^ Tfol, CD4^+^ Foxp3^+^ regulatory cells, and combinations of both. All subsequent analyses were conducted using R (version 4.3). Fluorescence intensity of all cells was transformed using the logicle method, and markers used solely for gating (Viability, CD3, CD19, CD4) were discarded. UMAP dimensionality reduction was performed using the uwot R package (https://cran.r-project.org/web/packages/uwot). Clustering was based on marker fluorescence intensity using the k-means algorithm, and resulting clusters were refined to reduce the final number via hierarchical-based metaclustering and supervised annotation. Heatmaps were generated with median expression values using the ComplexHeatmap R package (https://bioconductor.org/packages/release/bioc/html/ComplexHeatmap.html). Multidimensional scaling (MDS) and principal component analysis (PCA) were computed based on marker median expression values using the MASS R package (https://cran.r-project.org/web/packages/MASS) and the stats R package (https://stat.ethz.ch/R-manual/R-devel/library/stats).

### Sorting of live lymphocytes from cultured tonsillar whole cell suspensions

Following 24-hour *in vitro* incubation of tonsillar cell suspensions under stimulation alone or left untreated, the cells were stained for viability using 405/520 Viobility (1/500, Miltenyi). Subsequently, the cells were washed with 1X PBS and 5% FBS. Sorting of live lymphocytes (LiveDead^-^ FSC-A^lo^ SSC-A^lo^) was performed using a BD FACSAria II sorter (BD Biosciences).

### Single-cell RNA library construction and sequencing

Single-cell RNA library construction and sequencing were performed following the sorting of total live lymphocytes by flow cytometry: a single-cell library construction was performed using the Chromium Next GEM Single Cell 5’ Library and Gel Bead Kit v1.1 (PN-1000165), Chromium Next GEM Chip G (PN-1000127), Chromium Single-Cell 5’ Library Construction Kit (16 reactions) (PN-1000020), and Single Index Kit T Set A (96 reactions) (PN-1000123), according to the manufacturer’s protocol. Single-cell suspensions obtained from 20,000 cells, with barcoded gel beads and partitioning oil, were loaded onto Chromium Next GEM Chip G to generate single-cell bead-in-emulsion. Full-length cDNA along with cell barcode identifiers were then PCR-amplified to generate 5’ Gene Expression (GEX) libraries. These libraries were sequenced using a NovaSeq 6000 (Illumina) to achieve a minimum of 23,000 paired-end reads per cell for GEX. Reads were subsequently aligned to the GRCh38 human reference genome using Cell Ranger (v6.1.1, https://support.10xgenomics.com/single-cell-gene-expression/software/pipelines/latest/installation).

### Analysis of single-cell RNA-seq

The expression matrices resulting from reads alignment were analyzed using the Seurat R package^34^. Only high-quality cells were selected for subsequent analysis. The criteria for high-quality cells included a mitochondrial gene percentage below 10% and a number of unique molecular identifiers (UMI) between 200 and 15,000. Regularized negative binomial regression, based on the 3,000 most variable genes, was used to normalize the count matrix. Dimensionality reduction was performed using principal component analysis (PCA) based on the 3,000 most variable genes. Tfol were isolated by selecting the cells matching the following rules for the number of reads: *CD3* > 0, *CD4* > 0, *CXCR5* > 0, *CD19* = 0. Differential gene expression analysis was conducted on Tfol using the Wald t-test with p-values corrected using the false discovery rate (FDR) procedure.

### Study design and blood sample management of the Trans*i*mmunom cohort

Autoimmune and autoinflammatory patients as well as healthy volunteers were enrolled in the Trans*i*mmunom observational trial (NCT02466217)^35^. In our study, we analyzed patient samples from cohorts with at least 10 individuals: rheumatoid arthritis (n = 90), type 1 diabetes (n = 59), ankylosing spondylitis (n = 56), osteoarthritis (n = 47), Behçet disease (n = 38), systemic lupus erythematosus (n = 33), type 2 diabetes (n = 30), antiphospholipid syndrome (n = 23), Takayasu arteritis (n = 22), Sjögren’s syndrome (n = 19), granulomatosis with polyangiitis (n = 14), Crohn’s disease (n = 10), and healthy volunteers (n = 57). Blood samples were collected from all individuals to undergo multiple assays, performed by the Clinical Investigation Center for Biotherapies. Briefly, sera were used to quantify autoantibodies using enzyme-linked immunosorbent assay (ELISA), including antinuclear antibodies (anti-SSA52, anti-SSA60, anti-SSB, anti-Sm, anti-RNP, anti-Ro52, anti-nuclear DNA, anti-PmScl), anti-Ku, anti-TIF1, anti-cardiolipin IgG and IgM, anti-B2GP1 IgG and IgM, anti-Saccharomyces cerevisiae IgG and IgA, ANCA, anti-MPO, anti-PR3, anti-CCP, Latex, Waaler Rose, and anti-MBG. Cytokines and soluble receptors related to Tfh and Tfr activation, including IL-1β, IL-10, IL-2, sIL-1R1, and sIL-1R2, were quantified in the sera using multiplex assays (MILLIPLEX MAP Human Cytokine/Chemokine Growth Factor Panel A and Human Soluble Cytokine Receptor kits) with a MAGPIX analyzer according to the manufacturers’ instructions. Flow cytometry, performed with the Duraclone technology (Beckman Coulter) as described in Pitoiset *et al.*^36^, enabled the identification and quantification of cell populations^37,38^, among which only Tfh and Tfr were considered.

### Statistical analysis

After normality testing by Shapiro-Wilk test, P-values were calculated with Wilcoxon test for paired data and Mann-Whitney test for unpaired data. The statistical over-representation of gene list overlap with specific signatures was estimated using the hypergeometric test. Statistics provided in the results section are presented as median ± (1.58×IQR)/√n, where IQR represents the interquartile range and n the sample size.

## Results

### Human secondary lymphoid organs differ in their follicular T-cell composition and phenotype

To characterize Tfh and Tfr among major CD4^+^ T cell subsets from different secondary lymphoid organs (SLO), we collected 8 pediatric tonsils, as well as 8 paired spleens and mesenteric lymph nodes (mLNs) from unrelated healthy donors (**Figure 1A**). We hypothesized that these tissues would cover a wide range of Tfh and Tfr activation states, since they represent 3 types of SLO: mucosa-associated, blood-draining, and lymph-draining organs, respectively. Using flow cytometry, we identified 4 major CD4^+^ T cell subsets based on CXCR5 and Foxp3 expression: Tfh and Tfr within Tfol, Tconv, and Treg within non-Tfol (**Figure S1A**).

**Figure 1.**
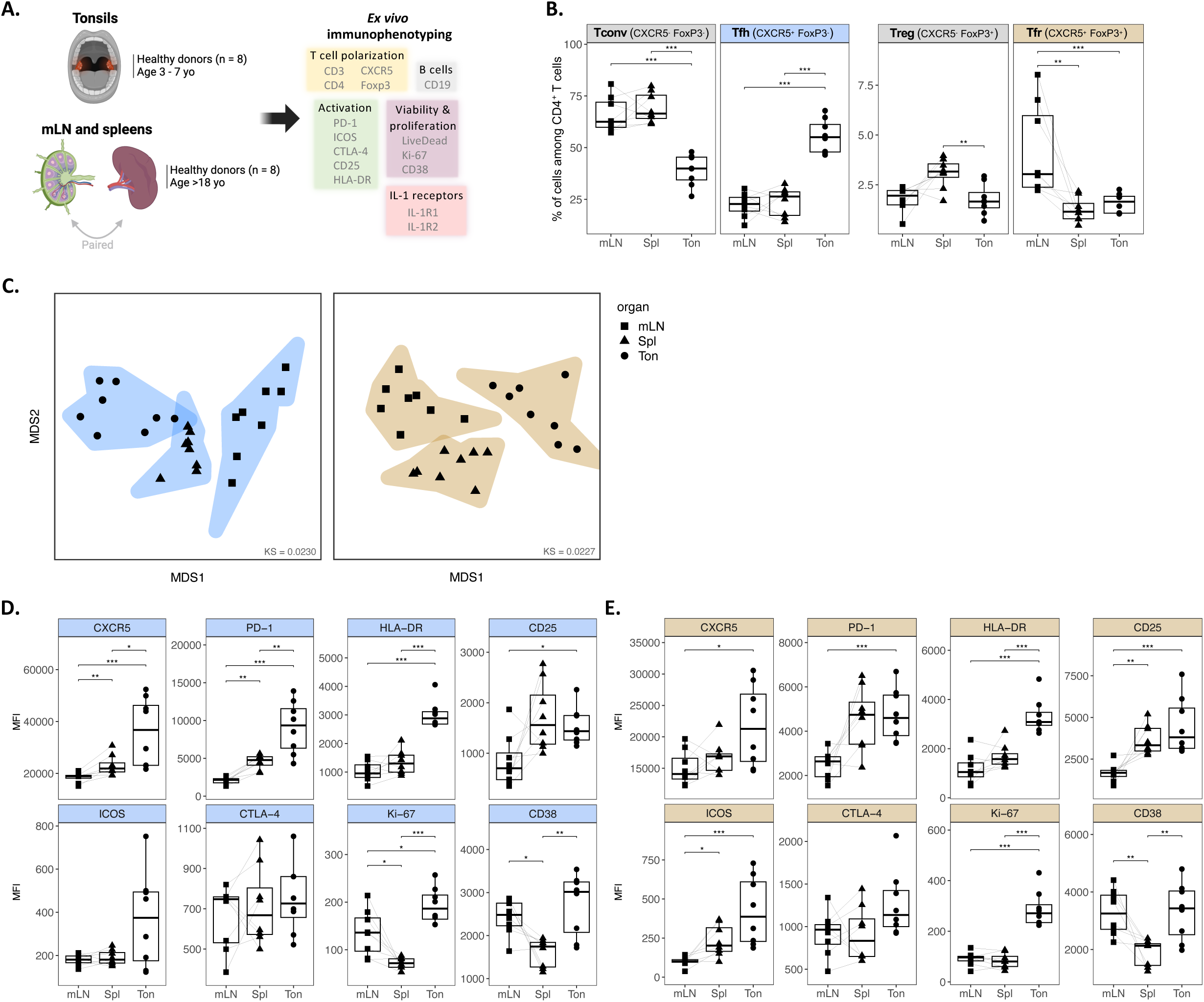
The abundance and phenotype of Tfol vary upon secondary lymphoid organs. (A) Experimental design for the immunophenotyping of cell suspensions extracted from healthy pediatric tonsils (n = 8), paired mesenteric lymph nodes (mLN, n = 8), and spleens (n = 8) from adult healthy donors. (B) Percentages of effector CXCR5^-^ Foxp3^-^ (Tconv) and CXCR5^+^ Foxp3^-^ (Tfh) cells, and regulatory CXCR5^-^ Foxp3^+^ (Treg) and CXCR5^+^ Foxp3^+^ (Tfr) cells among CD4^+^ T cells across lymphoid organs (n = 8 each). (C) Multidimensional scaling (MDS) plot displaying the similarities and dissimilarities between samples based on the MFI of all markers within Tfh (left, blue) and Tfr (right, gold) per sample, grouped by organ. (D-E) Mean fluorescence intensity (MFI) of markers of interest within Tfh (D) and Tfr (E) across lymphoid organs (n = 8 each). KS: Kruskal’s Stress. *P < 0.05; **P < 0.01; ***P < 0.001 by Mann-Whitney or paired Wilcoxon test.

The *ex vivo* abundance of Tfh among CD4^+^ T cells was significantly higher in tonsils than in spleen and mLNs (P < 0.001), as opposed to Tconv abundance which was decreased in tonsils (P < 0.001) (**Figure 1B**). The abundance of Treg differed in spleens with significantly increased frequency compared to tonsils (P < 0.01), while Tfr frequency was enriched in mLNs compared to spleens (P < 0.01) and tonsils (P < 0.001).

To address differences in Tfh and Tfr phenotypes between lymphoid organs, we integrated the expression of all markers by performing multidimensional scaling (MDS) on each population (**Figure 1C**). In Tfh (blue) and Tfr (brown), samples clustered separately and distinctly according to their organ of origin. Tfh and Tfr formed a gradient along the MDS1 dimension from tonsils to mLNs, aligning with the gradient of activation marker expressions.

We examined the phenotype of Tfh and Tfr across lymphoid organs by comparing the mean fluorescence intensity (MFI) of 8 markers related to cell activation (CXCR5, PD-1, HLA-DR, CD25, ICOS), proliferation (Ki-67, CD38), and function (CTLA-4) (**Figure S1B**). In Tfh, we observed a significant and gradual increase in the MFI of CXCR5, PD-1, HLA-DR, and CD25 within spleens and tonsils, compared to mLNs. ICOS and CTLA-4 were also increased following the same pattern, though not reaching significance (**Figure 1D**). Interestingly, both Ki-67 and CD38 MFI were significantly higher in tonsil Tfh, whereas their expression was downregulated in the spleen compared to paired mLNs, suggesting a lack of proliferating splenic Tfh. Tfh heterogeneity was observed regarding bimodal expression patterns of CXCR5, PD-1, and CD25, indicating variations in Tfh activation between organs and between donors (**Figure S1C**).

Similarly, Tfr displayed a significant and gradual increase in the MFI of activation markers (CXCR5, PD-1, HLA-DR, CD25, ICOS) in tonsils (**Figure 1E**), contrasting with its lower abundance in this organ. Ki-67 and CD38 expressions mirrored the ones of Tfh, underlining the higher proliferative phenotype of tonsillar Tfh and Tfr compared to splenic counterparts. The Tfr population exhibited heterogeneity regarding the expression pattern of CXCR5, Foxp3, PD-1, CD25, and CTLA-4, indicating distinct phenotypes of Tfr within lymphoid organs (**Figure S1C**).

In summary, our results show that the frequencies and phenotypes of Tfh and Tfr vary according to their SLO origin, suggesting a potential imprint of SLO on follicular T cell biology and highlighting pediatric tonsils as the SLO with the most activated Tfol.

### The expression of IL-1 receptors differs between Tfh and Tfr according to lymphoid cell activation

To investigate whether IL-1Rs contribute to the differential activation status of Tfol between lymphoid organs *ex vivo*, we first quantified the proportion of IL-1R1-expressing and IL-1R2-expressing cells among CD4^+^ T cells using flow cytometry (**Figures S2A and S2B**). Remarkably, the percentage of IL-1R1^+^ cells was significantly higher in tonsils, regardless of the CD4^+^ T cell subset, suggesting a link between increased tonsillar activation (**Figures 1C and 1D**) and IL-1R1 expression (**Figure 2A**). Notably, IL-1R1 expression was preferentially observed in regulatory subsets including Treg and Tfr, within all lymphoid organs (**Figure S2C**, top). Similarly, the MFI of IL-1R1 among IL-1R1^+^ Tfh and Tfr is enhanced in tonsillar Tfr compared to mLN counterparts (P < 0.05) (**Figure 2B**).

**Figure 2.**
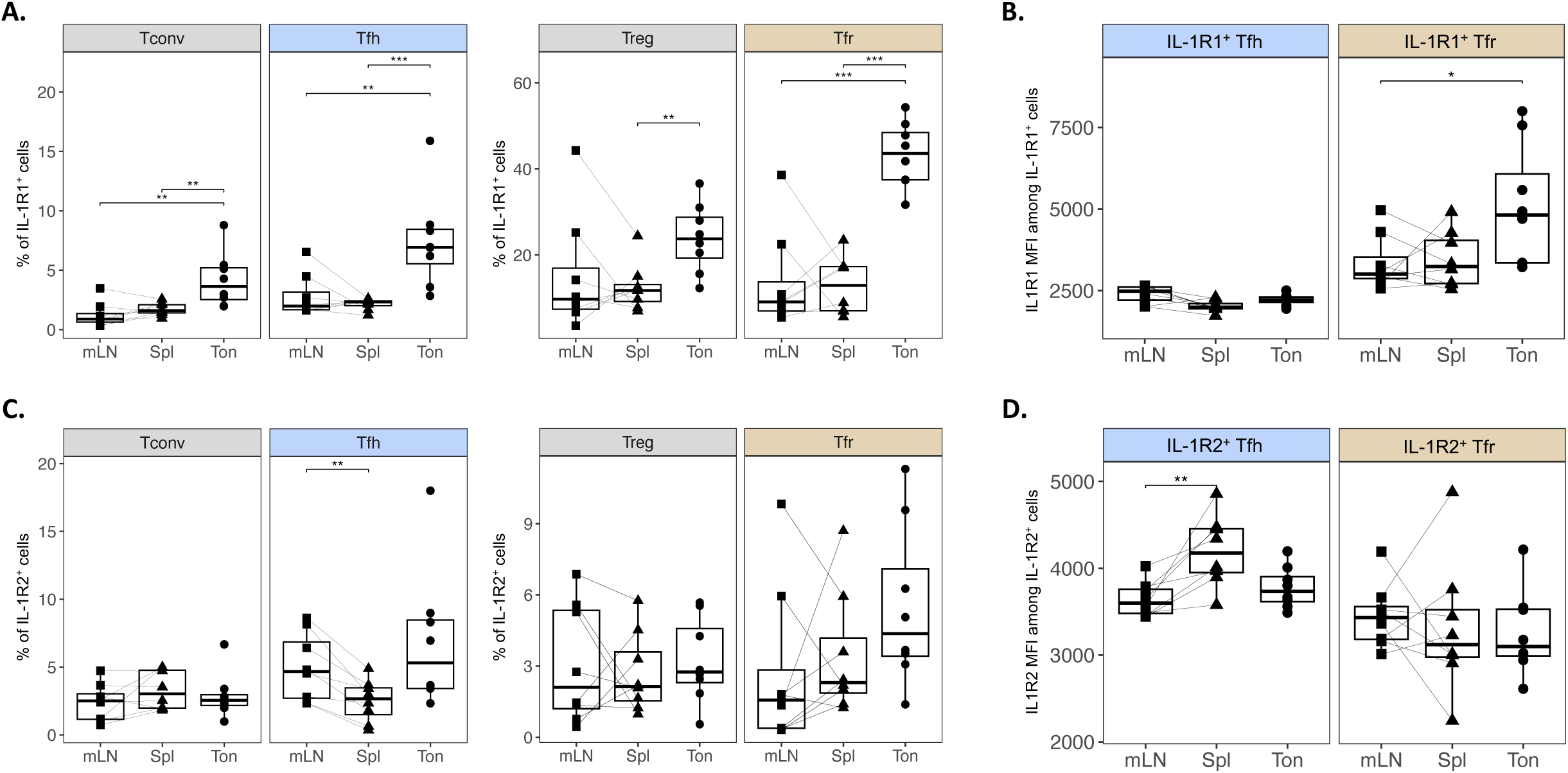
Tfh and Tfr display distinct patterns of IL-1Rs expression across secondary lymphoid organs. (A) Percentages of IL-1R1^+^ cells among CD4^+^ T cell subsets (n = 8 each). (B) Mean fluorescence intensity (MFI) of IL-1R1 within IL-1R1^+^ Tfh (left, blue) and Tfr (right, gold) across lymphoid organs (n = 8 each). (C) Percentages of IL-1R2^+^ cells among CD4^+^ T cell subsets (n = 8 each). (D) MFI of IL-1R2 within IL-1R2^+^ Tfh (left, blue) and Tfr (right, gold) across lymphoid organs (n = 8 each). **P < 0.01; ***P < 0.001 by Mann-Whitney or paired Wilcoxon test.

The proportion of IL-1R2^+^ cells among CD4^+^ T cell subsets varied to a lesser extent between organs (**Figure 2C**). However, we did observe that IL-1R2^+^ Tfh from mLNs (4.7 ± 2.3%) were significantly more abundant than in the spleens (2.7 ± 1.1%) (P < 0.01). Thus, in Tfh, the expression of the IL-1 antagonist receptor matches that of proliferation markers Ki-67 and CD38 (**Figure 1D**). Of note, the proportion of cells expressing IL-1R2 was higher among Tfh than Tconv subsets in tonsils and mLNs (**Figure S2C**, bottom). The MFI of IL-1R2 solely differed within IL-1R2^+^ Tfh, not Tfr counterparts, the former being more elevated within spleens than mLNs (P < 0.01) (**Figure 2D**). Interestingly, the proportion of IL-1R2^+^ Tfh appeared dissociated from the MFI suggesting that a significant proportion of Tfh cells in mLNs express IL-1R2 weakly.

When comparing patterns of IL-1R1 and IL-1R2 expression between Tfh and Tfr across organs, Tfr cells, and not Tfh cells, seemingly displayed a multimodal expression of IL-1R1 with emphasis on tonsils (**Figure S2D**). Such heterogeneity indicates that multiple Tfr subsets are found according to IL-1R1 expression and lymphoid activation.

Overall, the varying patterns of IL-1R1 and IL-1R2 expression between organs underscore distinct phenotypes following the activation status of Tfh and Tfr cells.

### The expression of IL-1Rs captures a distinct, activated Tfr population and is associated with Tfh maturation

We hypothesized that the expression of IL-1Rs could contribute to the different stages of follicular T-cell activation. To assess this contribution, we merged flow cytometry data from tonsils, spleens, and mLN to set up a metaorgan with the aim of comparing Tfh and Tfr activation stages according to their organ of origin. We focused our analysis on Tfr (CD4^+^ CXCR5^+^ Foxp3^+^), Tfh (CD4^+^ CXCR5^+^ Foxp3^-^) and Treg (CD4^+^ CXCR5^-^ Foxp3^+^), all three being the populations with higher IL-1R1 or IL-1R2 expression.

To study the phenotype of the cells involved in our metaorgan, we used unsupervised methods which offer the advantage of unbiasedly studying cell populations with multiple markers. Using the *ex vivo* expression of 11 markers among the cells of the metaorgan, we performed UMAP dimensionality reduction (**Figure S3A**). Then, we identified clusters of interest (**Figure 3A**) based on an analytical strategy encompassing unsupervised clustering, metaclustering, and literature-based annotation (**Figure S3B**). This allowed the identification of 5 clusters that discriminated cells upon their activation status: two Tfh clusters with increasing CXCR5 and PD-1 expression (namely *Tfh* and *GC-Tfh*), two Tfr clusters that differed in the expression of Ki-67, HLA-DR, ICOS, and CTLA-4 (namely *resting Tfr* and *activated Tfr*), and one Treg cluster (**Figure 3B**). *Activated Tfr* (*act. Tfr*) phenotypically clustered together with *GC-Tfh* as a result of a shared expression of PD-1, Ki-67, HLA-DR, ICOS, and CD38, while the expressions of CD25, Foxp3, and CTLA-4 markers were rather shared with *resting Tfr* (*rest. Tfr*) and Treg clusters. The abundance of each cluster between SLO revealed that the two Tfh clusters are not equally present, since *GC-Tfh* had higher frequencies in activated tonsils versus mLNs and spleens (P < 0.001) (**Figure S3C**). This confirms that tonsils display activated Tfol (**Figure 1E**) and attests to the validity of the metaorgan clustering approach. In contrast, all regulatory cell clusters had significantly lower frequencies within tonsils (**Figure S3C**).

**Figure 3.**
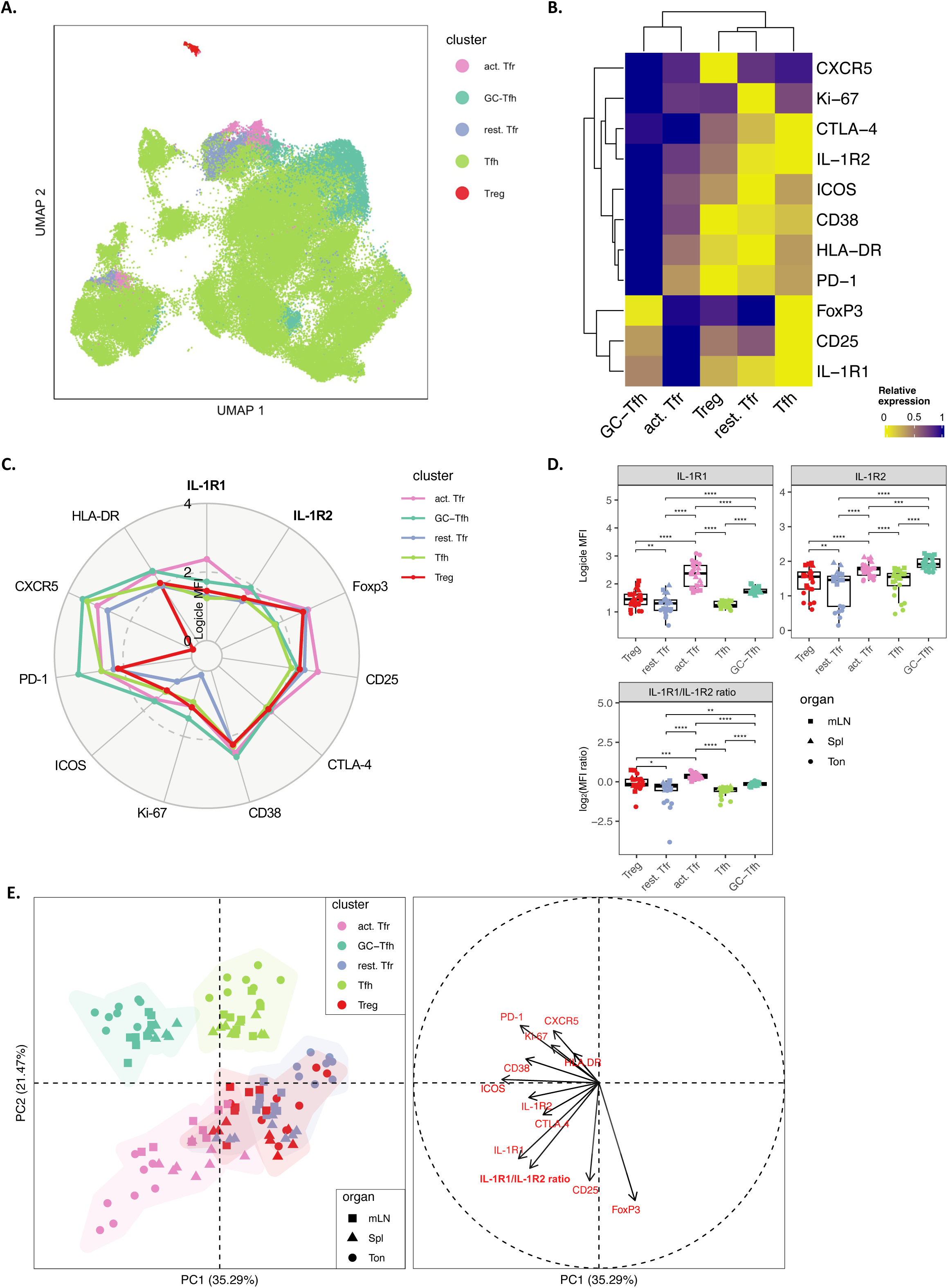
IL-1R1 and IL-1R2 differential expression defines an activated Tfr population and shapes Tfh maturation. (A) UMAP representation of CD4^+^ CXCR5^+^ (Tfh and Tfr) cells and CXCR5^-^ Foxp3^+^ cells (Treg) across all secondary lymphoid organs (n = 24), colored by identified clusters of cells. (B) Heatmap showing relative marker expression of the five identified cell clusters. (C) Radar plot displaying median logicle-MFI of markers of interest across the five clusters. (D) Logicle-transformed MFI of IL-1R1 and IL-1R2 (top) and IL-1R1 MFI ratios (bottom) across the five clusters within all lymphoid organs (n = 24). (E) Principal component analysis (PCA) displaying the separation of samples accordingly to the five identified cell clusters within all lymphoid organs (left) and its associated correlation circle showing the contribution of each marker to the principal components indicated with vectors (right). *P < 0.05; **P < 0.01; ***P < 0.001; ****P < 0.0001 by Mann-Whitney test.

The quantification of marker expression, summarized as a radar plot of logicle-transformed MFI per cluster, highlighted the activated phenotype of *GC-Tfh* compared to *Tfh*, regarding ICOS, HLA-DR, CD25, CD38, CTLA-4, and Ki-67 (**Figure 3C**, **Figure S3D**). Similarly, *activated Tfr* displayed an upregulation of these activation and proliferation markers relatively to *resting Tfr* and *Treg*. In detail, IL-1R1 logicle-MFI is significantly higher in *act. Tfr* (P < 0.0001), with an even higher expression in tonsils (**Figure 3D**). *GC-Tfh* also displayed an increased IL-1R1 expression compared to *Tfh* (P < 0.0001) (**Figure 3D**). Accordingly, these activated clusters also upregulated IL-1R2 expression compared to their non-activated counterparts (P < 0.0001) (**Figure 3D**). When performing the ratio of surface expression of IL-1R1/IL-1R2 MFIs, which assesses the responsiveness of cells to IL-1β, *act. Tfr* and *GC-Tfh* also displayed a significant increase compared to their resting counterparts (P < 0.0001). Of note, solely *act. Tfr* had a positive median log_2_(MFI ratio), thus indicating a more important sensitivity to IL-1β, while *rest. Tfr* are less likely to respond to IL-1β (**Figure 3D**). Therefore, IL-1Rs appear as surrogate markers of Tfh and Tfr activation.

To apprehend the weight of all markers to the clustering of the clusters across organs, we submitted our 5 clusters to a principal component analysis (PCA) (**Figure 3E**). The PCA well separated Tfh from Tfr and Tregs, these latter being intermingled. PCA points towards a continuum from *Tfh* to *GC-Tfh* following CXCR5 and PD-1 expression, with tonsillar samples being the most driven by this activated phenotype compared to spleens and mLNs (**Figure 3E**). *Rest. Tfr* strongly overlapped with *Treg* cluster, indicating their phenotypical similarity. Strikingly, *act. Tfr* cluster appeared more segregated with a marked distinction of tonsillar samples (**Figure 3E**), due to the contribution of IL-1R1 and the IL-1R1/IL-1R2 ratio. *Act.Tfr* share markers with regulatory clusters (CTLA-4, CD25, Foxp3) and co-stimulation (ICOS). IL-1R2, on the other hand, partly drives the *GC-Tfh* cluster, suggesting that IL-1R2 expression on *GC-Tfh* contributes to the maintenance of a GC-like phenotype, while IL-1R1 rather orientates to a Tfr phenotype. CD38 and ICOS, while contributing to *GC-Tfh* cluster, are also shared with *act. Tfr* and appear as markers of follicular T activation (**Figure 3E**).

Taken together, our data indicate that the Tfr population is heterogeneous, one subset being activated in accordance with the expression of IL-1R1 over IL-1R2. While Tfh activation is also associated with increased IL-1R1 expression, IL-1R2 seems critical for GC-Tfh maintenance by potentially modulating the sensitivity of the cells to IL-1β signaling.

### IL-1β-responsive Tfr exhibit dual suppressive and effector functions similar to Treg and Tfh

To decipher whether differential expression of IL-1Rs regulates Tfh and Tfr functions, we analyzed tonsils (n = 4) and spleens (n = 3) given the higher expression levels of IL-1Rs among Tfh and Tfr subsets. Whole-lymphoid cell suspensions were immunophenotyped using an intracellular flow cytometry panel comprising IL-2, IL-4, IL-10, and IL-21 cytokines **(Figure 4A** and **Figure S4A)**. We performed a joint unsupervised analysis of Tfh, Tfr, and Treg cells based on the expression of 9 markers (**Figure S4B** and **Figure 4B**). We identified the 5 same clusters based on the previous annotation strategy comprising the core markers CXCR5, Foxp3, PD-1, and Ki-67 **Figure 4C**). *Act. Tfr* and *Treg* clustered together due to similar IL-10, IL-21, and Ki-67 expression patterns, whereas *Tfh* and *rest. Tfr* clustered separately because of very poor cytokine and activation marker expression profiles (**Figure 4C**). *GC-Tfh* clustered apart with a unique relative expression of IL-4, CXCR5, and PD-1.

**Figure 4.**
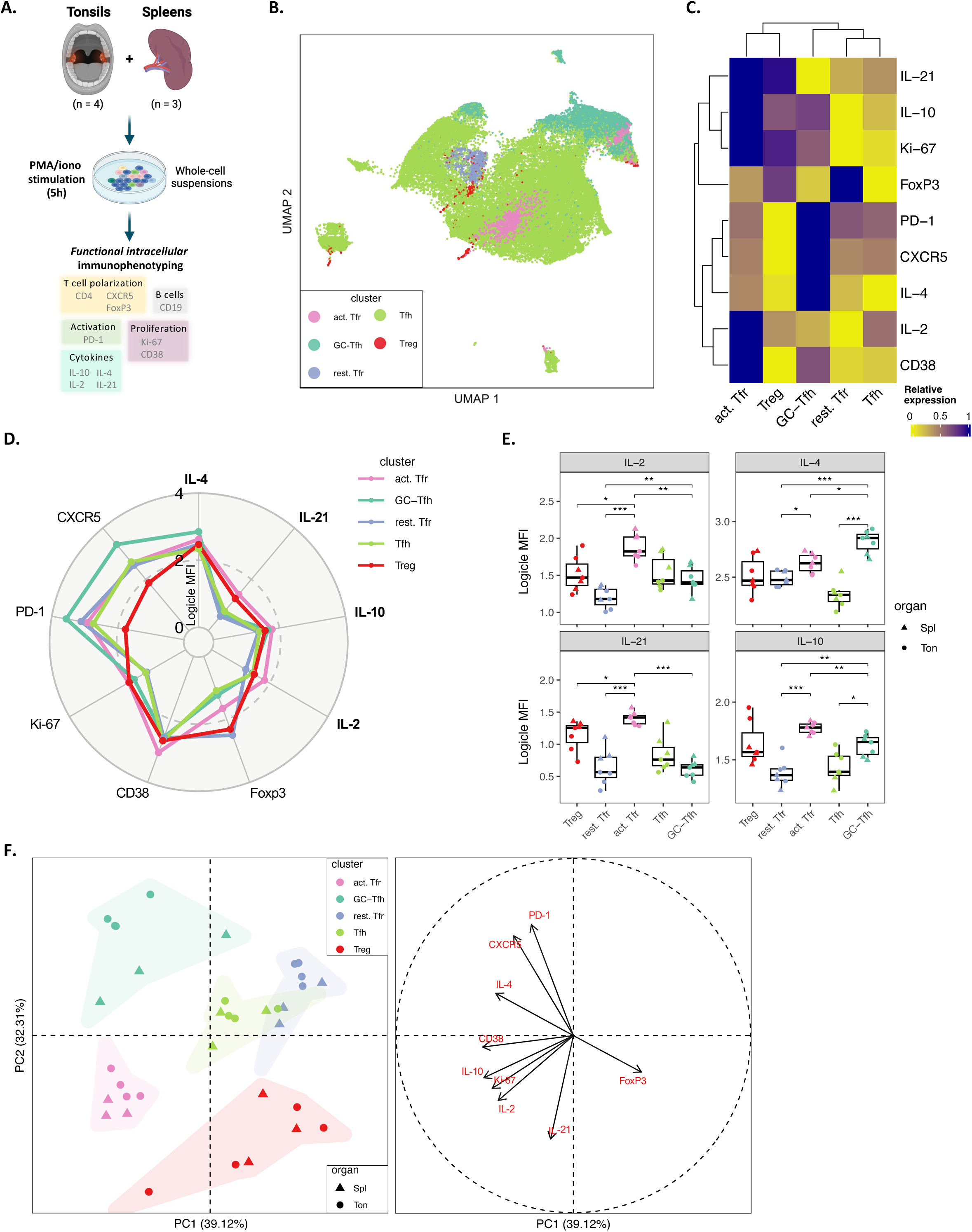
Activated Tfr with high IL-1R1/IL-1R2 ratio are functionally similar to both Treg and Tfh. (A) Experimental design of in vitro lymphoid cell stimulation from selected tonsils (n = 4) and spleens (n = 3) for functional immunophenotyping using flow cytometry. (B) UMAP representation of CD4^+^ CXCR5^+^ (Tfh and Tfr) cells and CXCR5^-^ Foxp3^+^ cells (Treg) across selected secondary lymphoid organs (n = 7), colored by identified clusters of cells. (C) Heatmap showing relative marker expression of the 5 identified cell clusters. (D) Radar plot displaying median logicle-MFI of 9 markers of interest across the 5 clusters. (E) Logicle-transformed MFI of IL-2, IL-4, IL-21 and IL-10 across the 5 clusters. (F) Principal component analysis (PCA) displaying the separation of samples accordingly to the five identified cell clusters within all lymphoid organs (left) and its associated correlation circle showing the contribution of each marker to principal components indicated with arrows (right). *P < 0.05; **P < 0.01; ***P < 0.001 by Mann-Whitney test.

In detail, the absolute logicle-transformed MFI within the radar plot indicated that the *act. Tfr* cluster produces the most amounts of cytokines and is the most proliferative based on CD38 and Ki-67 markers (**Figure 4D**). In contrast, *rest. Tfr* and *Tfh* are the clusters with weaker cytokine production and proliferation status. Indeed, the *act. Tfr* cluster displayed higher expression of IL-2 (P < 0.001), IL-4 (P < 0.05), IL-21 (P < 0.001), and IL-10 (P < 0.001) than *rest. Tfr*, and higher expression of IL-2 (P < 0.01), IL-10 (P < 0.01), and IL-21 (P < 0.001) than *GC-Tfh* (**Figure 4E**). The latter only showing increased expression of IL-4 (P < 0.05) relative to *act. Tfr* (**Figure 4E**).

Finally, we performed a PCA analysis to determine the contribution of each marker to the functional phenotype of the identified clusters (**Figure 4F**). While the *GC-Tfh* cluster is strongly driven by CXCR5, PD-1, and IL-4 expression, the *act. Tfr* cluster is driven by IL-10, IL-2, and Ki-67 expression. Both clusters share the contribution of CD38 to their phenotype, and the *act. Tfr* cluster also shares strong IL-21 and Foxp3 expression with Treg thus appearing as a cluster that is functionally halfway between *GC-Tfh* and *Treg*.

In sum, *act. Tfr* cells are functionally close to both GC-Tfh subsets and Treg cells in terms of cytokine expression. Combined with IL-1Rs expression, the cytokine expression analysis discriminates activated from resting states of Tfol.

### IL-1β responsiveness is finely tuned and potentiates Tfh and Tfr activation

To investigate how Tfh and Tfr responsiveness to IL-1β is regulated during cell activation, we compared the expression of IL-1Rs between stimulated (stim) and non-stimulated (no stim) Tfh and Tfr. We cultured total tonsillar lymphoid cells (n = 8) for 24 hours and stimulated T cells with anti-CD3/CD28 beads alone (stim) or with IL-1β (stim+IL-1β) or with IL-1R1 inhibitor Anakinra (stim+Anakinra) (**Figure 5A**).

**Figure 5.**
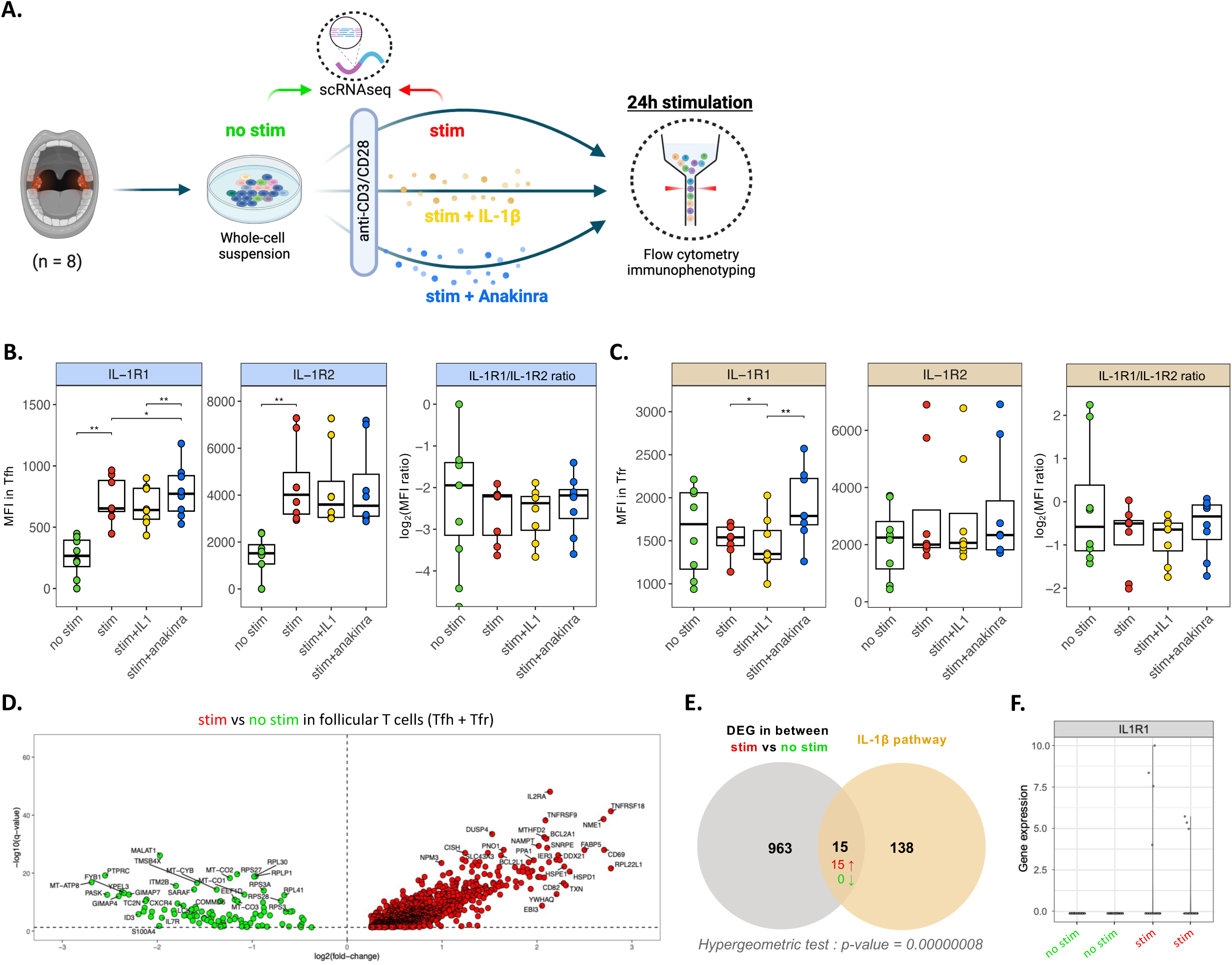
IL-1β potentiates tonsillar Tfh and Tfr activation in vitro. (A) Experimental design for whole-cell tonsillar suspensions (n = 8) stimulated with anti-CD3/CD28 beads for 24 hours, either in the presence of IL-1β (0.5 µg/ml), Anakinra (1 µg/ml), or alone. The expression of IL-1R1 and IL-1R2 was measured using flow cytometry. (B-C) Boxplots showing the MFI of IL-1R1 and IL-1R2, and their ratio among Tfh (B) and Tfr (C) from whole-cell tonsillar cultures compared to no stimulation (n = 8). (D) Volcano plot showing differentially expressed genes (DEG) in stimulated *CXCR5*^+^ Tfol relative to no stimulation, analyzed using single-cell RNA sequencing (n = 2 tonsils). (E) Venn diagram displaying the overlap between DEG and the genes belonging to IL-1β pathway obtained from Wikipathway. (F) Expression of the *IL1R1* that was one of the DEG overlapping with the IL-1β pathway. *P < 0.05; **P < 0.01 by paired Wilcoxon test.

First, anti-CD3/28 stimulated Tfh displayed enhanced IL-1R1 and IL-1R2 MFI (**Figure 5B**). Adding IL-1β did not significantly change IL-1R1 or IL-1R2 expression levels. However, inhibiting endogenous IL-1β signaling with Anakinra increased IL-1R1 expression among Tfh (P < 0.05). This suggests that IL-1β is produced *in vitro* during stimulation, potentiated by antigen-presenting cells. Accordingly, the frequency of IL-1R1^+^ and IL-1R2^+^ Tfh followed the same patterns (**Figure S5A**). Of note, the ratio of IL-1R1/IL-1R2 MFIs remained unchanged in all conditions, indicating that sensitivity to IL-1β was maintained after stimulation (**Figure 5B**).

In contrast with Tfh, Tfr did not exhibit significant IL-1Rs variations upon stimulation, potentially due to an already high expression at steady state (**Figure 5C**). However, IL-1R1 MFI in Tfr was significantly diminished with the addition of IL-1β compared to control (P < 0.05), and Anakinra restored normal amounts compared to the stim+IL-1β (P < 0.01). However, IL-1R2 expression did not vary in Tfr. The frequency of IL-1R1^+^ and IL-1R2^+^ Tfr matched the levels of expression indicated by MFI (**Figure S5B**). Similar results were observed on isolated Tfh and Tfr stimulated with IL-1β (not shown). The sensitivity of Tfr to IL-1β was maintained regarding the stability of the IL-1R1/IL-1R2 MFI ratio after stimulation (**Figure 5C**). Regarding cell abundance, the Tfr/Tfh ratio did not significantly vary between conditions, although the addition of Anakinra tended to decrease the ratio, and thus unfavoured Tfr (**Figure S5C**).

To confirm that follicular T-cell activation was indeed associated with IL-1β signaling, we performed single-cell RNA sequencing on live tonsillar lymphoid cells in stim and no stim conditions (n = 2) (**Figure 5A**). We selected *CD4* and *CXCR5*-expressing T cells to perform dimensionality reduction followed by differential gene expression analysis (**Figure 5D**). The volcano plot features the magnitude of gene expression expressed as a fold change (log_2_-transformed) and the statistical evaluation via the q-value (-log_10_-transformed). The latter highlighted several upregulated genes during stimulation, whereas fewer genes were shown to be downregulated. Genes involved in T cell proliferation and early activation^39^ such as *IL2RA* (CD25), *CD69*, *TNFRSF9* (4-1BB), *TNFRSF18* (GITR) and *NME1* were found upregulated. On the contrary, downregulated genes comprised *IL7R* (CD127) and *S100A4* which are associated to a resting T cell phenotype, as well as *ITM2B* known to be repressed by *BCL6*^40^. The functional enrichment analysis using genes upregulated during stimulation revealed a significant enrichment of pathways linked to T cell proliferation and function such as T cell receptor signaling and apoptosis, IL-2, and NF-κB pathways (P < 0.0001).

Then we assessed whether DEG overlapped with genes from IL-1β pathway. We constituted an exhaustive 153-gene list of the pathway by searching public databases (WikiPathways, Reactome, KEGG pathway) for all genes involved in sensing, processing, signaling, and regulation of IL-1β and its direct targets (**Table 1**). Among the 153 genes composing the pathway, 15 were significantly found (P < 0.0001) among DEG, all of them being upregulated (**Figure 5D**). Interestingly, *IL1R1* expression significantly increased during stimulation (**Figure 5F**), together with *NFKB1*, a key downstream mediator of IL-1β signaling. By using a curated public IL-1β pathway (*WP4496*, WikiPathways), we found 5 of our DEG significantly upregulated among the 33 genes of the pathway (P = 0.000427) (**Figure S5D**).

Taken together, our transcriptomic results underline the ability of IL-1β to support Tfh and Tfr activation through NF-κB signaling. In response to IL-1β exposure, a decrease of IL-1R1-expressing cells is observed, while inhibition of IL-1β induces an augmented expression of IL-1R1.

### Increased peripheral IL-1β and cTfr as surrogate markers of autoantibody production in autoimmune patients

Since IL-1β affects the activation and function of Tfol *in vitro*, we explored whether the production of different types of autoantibodies could be linked to disrupted Tfh and Tfr balance and elevated IL-1β levels. To address this question, we analyzed flow cytometry immunophenotyping, cytokine and autoantibody detections from blood samples of a cohort of patients with various autoimmune and autoinflammatory diseases including rheumatoid arthritis (n = 90), type 1 diabetes (n = 59), ankylosing spondylitis (n = 56), osteoarthritis (n = 47), Behçet disease (n = 38), systemic lupus erythematosus (n = 33), type 2 diabetes (n = 30), antiphospholipid syndrome (n = 23), Takayasu arteritis (n = 22), Sjögren’s syndrome (n = 19), granulomatosis with polyangiitis (n = 14), Crohn’s disease (n = 10), as well as healthy donors (n = 57).

We grouped individuals based on their autoantibody status and analyzed the levels of cTfr over cTfh cells (**Figure 6A**). While the percentages of cTfh and cTfr did not differ significantly between autoantibody-positive and -negative individuals, the cTfr/cTfh ratio was higher in those who were autoantibody-positive (**Figure 6A**). Among autoantibody-positive individuals, we quantified the number of different autoantibodies (from 1 to 8) and grouped those with three or more autoantibodies (≥3) (**Figure 6B**). We found that higher autoantibody diversity was associated with an increased cTfr/cTfh ratio. Notably, we also found a significant correlation (R = 0.55 and P < 0.001) between cTfr and cTfh across all individuals (**Figure S6A**).

**Figure 6.**
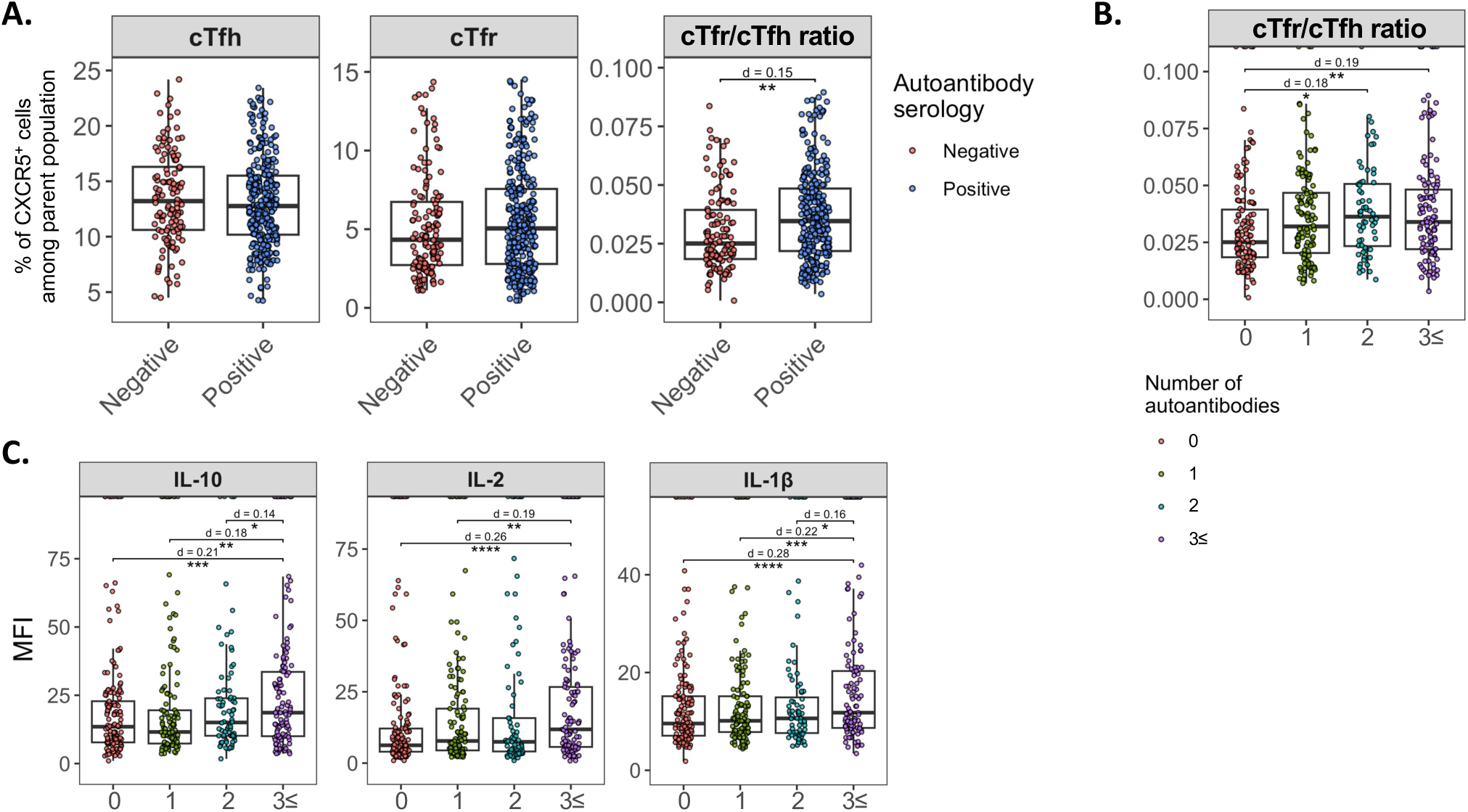
Higher autoantibody load is associated with increased cTfr and IL-1β-related molecules in the blood of autoimmune patients. (A) Frequency of circulating Tfh (cTfh) and circulating Tfr (cTfr) cells, as well as the cTfr/cTfh ratio, in relation to the autoantibody serology of healthy individuals and patients from the Trans*i*mmunom cohort. cTfh: CXCR5^+^ cells among CD3^+^ CD4^+^ T cells; cTfr: CXCR5^+^ cells among Foxp3^+^ CD127^-^ Treg. Negative: n = 152; positive: n = 327. (B) cTfr/cTfh ratio among all individuals according to their number of autoantibodies. (C) MFI measurements of peripheral cytokine levels according to the number of autoantibodies in all individuals. *P < 0.05; **P < 0.01; ***P < 0.001; ****P < 0.0001 by paired Wilcoxon test. d: Cliff’s delta effect size. cTfr: circulating Tfr; cTfh: circulating Tfh; sIL-1R1: soluble IL-1R1; sIL-1R2: soluble IL-1R2.

We also measured the levels of IL-10, IL-2, and IL-1β and found they were upregulated with higher autoantibody diversity (**Figure 6C**). Soluble IL-1R1 (sIL-1R1) levels were higher in individuals with more autoantibodies, while soluble IL-1R2 (sIL-1R2) levels remained unchanged (**Figure S6B**). This suggests that cells undergoing strong activation and stimulation by IL-1β release IL-1R1 as a regulatory response to inflammation.

Overall, our findings indicate that higher autoantibody diversity is associated with increased peripheral IL-1β and cTfr levels, and the release of sIL-1R1.

## Discussion

We describe the varying abundance and phenotypic heterogeneity of Tfol across 3 distinct lymphoid organs — tonsils, spleens, and mesenteric lymph nodes. Using *ex vivo* flow cytometry, we challenge the traditional static view of Tfh and Tfr by addressing their activation stages as a continuum, which is associated with increased expression of IL-1R1 at later stages. In Tfr cells, when IL-1R1 overcomes IL-1R2 expression, we describe an activated subset characterized by a shared phenotype and function with both Treg and GC-Tfh. Single-cell RNA sequencing and *in vitro* stimulation underscore that follicular T-cell activation requires IL-1β signaling through IL-1R1. As a result, Tfh and Tfr can vary the expression of IL-1R1 on their surface to modulate responsiveness to IL-1β. The exploration of IL-1β-related molecules on autoimmune individuals from the Trans*i*mmunom cohort revealed that IL-1β, circulating Tfr and IL-10 are surrogate markers of higher autoantibody load.

In summary, we show that Tfr heterogeneity across lymphoid organs is linked to IL-1R1 and IL-1R2 differential expression. Follicular T cell activation through IL-1β signaling could orchestrate the emergence of a Tfr subset with dual suppressive and helper functions. Excess of IL-1β during GC reaction may promote Tfr emergence and autoreactive antibody production.

Our work demonstrates that GC-Tfh, which are classically thought to be an end-stage of Tfh differentiation, appears as a potential seeder for the Tfr pool. As a result, two main Tfr subsets are found within human SLO: one is activated, phenotypically close to *GC-Tfh* and characterized by the expression of IL-1R1, CD25, CTLA-4, CD38, and Ki-67, as well as the production of IL-10, IL-2, IL-21, and IL-4, while the other Tfr subset is present at steady state, phenotypically closer to Treg and Tfh.

Although we did not experimentally assess trajectories leading to Tfr induction, a recent study demonstrates that GC-Tfh acquire CD25 and Foxp3 before differentiating into CD38^+^ CD25^+^ ICOS^+^ PD-1^+^ CTLA-4^+^ “iTfr”, located within the GC, clonally related to Tfh and able to promote B cell maturation via IL-10 production^22^. These observations are in line with the phenotype of our *act. Tfr* cluster, which shares properties with both *GC-Tfh* and *Treg.* Our findings extend this mechanism and suggest that IL-1β could drive *GC-Tfh*-to-*act. Tfr* transition. In contrast, it was shown that “nTfr” (CD38^-^) were clonally related to Treg and retained suppressive functions while remaining at the T-B border, indicating that our resting Tfr could emerge from the Treg lineage independently of IL-1β signaling.

The existence of a novel Tfr subset with helper properties challenges the current dogma. However, there is increasing evidence of Tfr having a dual helper and suppressive role in the context of GC reaction^41–43^. In particular, the role of CD25 has been debated over the past years given the discordance between murine studies, stating that GC-resident Tfr lose CD25 expression^29,44^, while human studies assert the maintenance of CD25 expression^22,45^. Le Coz *et al.* demonstrated that transition from *GC-Tfh* to *act. Tfr* phenotype is enhanced by IL-2 supplementation *in vitro*^22^. In line with those results in humans, we provide evidence that *act.Tfr* highly express CD25. We propose a model in which IL-1β also favors Tfr transition due to excess of IL-1R1 over IL-1R2 expression in Tfh, giving rise to a GC-resident, Tfh-derived Tfr population.

In our single-cell transcriptomics results, activated Tfol showed enhanced GC-like phenotype, associated with IL-1β/NF-κB signaling as well as IL-2 signaling. In fact, among the upregulated genes, we found the high affinity IL-2 receptor chain *IL2RA* (CD25): while IL-2 is known for its T cell stimulation effects, it also inhibits follicular T cell phenotype and thus carries a regulatory function during Tfh maturation^9^. Noteworthy, *CXCR4* was downregulated during T cell activation; its loss of expression is associated with preferential localization of Tfh in the GC light zone and distinguishes a Tfh phenotype with improved B cell help functions^46^. This downregulation among *GC-Tfh* was also observed *in vitro* following splenic T-cell stimulation^47^.

*In vitro* stimulated tonsillar T cells manifested a removal of surface IL-1R1 when IL-1β is added, which may occur in case of excessive IL-1 agonism. This mechanism can be achieved through IL-1R1 internalization^48^. Also, the receptor can be cleaved and secreted as a soluble form, thus sequestering extracellular IL-1β when present in high concentrations^49^. On the contrary, culture with Anakinra increased IL-1R1 surface expression.

Finally, we explored a cohort of individuals with distinct autoimmune and inflammatory conditions, as well as healthy volunteers, aiming at deciphering cTfh, cTfr, and IL-1β-related molecules. The presence of GC-related cytokines and circulating Tfol represent a biomarker of ongoing GC activity within tissues. In this study, we show that cTfh and cTfr amounts are correlated, and the cTfr/cTfh ratio increases with higher autoantibody diversity. Our data imply that IL-1β induced during inflammation may act on cTfr to support autoantibody production through IL-10 and Ki-67, potentially contributing to autoimmune pathogenesis. These results contrast with the dichotomic vision of Tfr being protective against autoimmunity and Tfh being detrimental by facilitating the production of autoantibodies. This can be due to the capacity of cTfr to promote the formation of ectopic lymphoid structures within inflammation sites, thus sustaining pathogenic GC reaction^23,50,51^. Accordingly, we show that peripheral IL-10 is found upregulated with higher autoantibody diversity, suggesting that Tfr-derived IL-10 can promote B cell maturation and antibody production^52^. Similarly, IL-1β levels followed the same pattern, indicating that cTfr increase can be attributed to the specific IL-1β-driven differentiation of *activated Tfr*. Alternatively, increased cTfr associated with autoantibodies may reflect an immune response aimed at unsuccessfully controlling the pathogenic mechanism.

The increase of sIL-1R1 in the blood of individuals with a high number of autoantibodies is in line with decreased surface IL-1R1 expression upon culture with IL-1β. The release of sIL-1R1 in the blood of individuals could be linked to excessive IL-1β binding and thus shedding of the receptor, serving as a biomarker of inflammation and autoantibody production. In contrast, decreased sIL-1R2 represents a lack of suppressive activity, which is correlated with joint destruction in arthritis^53^. Inflammation-induced IL-1β could promote the differentiation of a specific *act. Tfr* subset endowed with *GC-Tfh* functions.

Overall, our unique approach combining the phenotype of distinct lymphoid organs with different activation states has enabled us to characterize follicular T cell heterogeneity involving IL-1β-responsiveness, which appears to be a central cytokine for Tfh and Tfr modulation during human GC reaction.

### Limitations of the study

The present study has some limitations that must be considered when interpreting the results. One key limitation concerns the origin of the samples, either from healthy adult donors (spleen and mLNs) or from healthy pediatric donors (tonsils). As a result, Tfol from pediatric tissues may display a particular phenotype, different from the one of adult tonsillar Tfol. In addition, functional experiments for the characterization of cytokine production were conducted using only two types of organs: tonsils and spleen, which displayed the most and the least proliferative Tfol, respectively. This limited selection, while covering a wide range of activation states, may not fully represent the diversity of human Tfol activation features.

Our *in vitro* and scRNA-seq experiments were confined to tonsillar tissues, exhibiting high levels of Tfol activation and large amounts of cells. This limits our ability to generalize IL-1β-mediated activation patterns to tissues with lower levels of baseline activation. Expanding the scope to include tissues with varying activation states would provide a more comprehensive understanding of the broader applicability of IL-1β response profiles.

Another limitation is the lack of direct experimental evidence for Tfh-to-Tfr differentiation under the influence of IL-1β signaling. Although our results highlight IL-1β’s role in modulating Tfh and Tfr phenotypes and activation, further experiments are needed to provide concrete proof of this differentiation pathway, such as lineage tracing assays. In line with this limitation, our scRNA-seq approach did not allow the identification of *FOXP3*^+^ cells, which is essential for the identification of Tfr and could reveal TCR sharing between Tfh and Tfr. The work of Le Coz *et al*. demonstrated that Tfr indeed display TCR repertoire overlap with Tfh, indicating a common origin^22^. Yet, we managed to identify these Tfh-originated Tfr using the same markers, which strongly suggests the pivotal role of IL-1β in such differentiation.

An additional point of consideration is the endogenous production of IL-1β suggested in our *in vitro* experiments when Anakinra is added into culture. We could not assess the concentration of IL-1β nor soluble IL-1 receptors in culture supernatants, which would provide valuable insights into the mechanisms of regulation involved. Moreover, although the elevated IL-1β concentration employed in our experiments was instrumental in revealing underlying mechanisms, it may not fully replicate physiological conditions.

## Acknowledgements

We thank all donors and their families for providing pediatric tonsils, adult spleens, and mesenteric lymph nodes. We thank Dr. Eric Savier for providing adult organ samples from the digestive surgery department at Pitié-Salpêtrière Hospital. We thank Paul Stys for quality control of Trans*i*mmunom databases and Widad Nefida for optimization of protocols from the Immunology Immunopathology Immunotherapy research unit. We also thank the healthy individuals, patients, and all clinical staff members involved in the Transimmunom trial.

## Author contributions

Conceptualization: BB, DK, NT, PE, RJM, RV, SGD; Methodology: AB, GF, NT, RJM, RV, SGD; Experimentation: BG, LP, NC, RJM, RV, SAM; Data analysis: HV, JD, MB, HD, NTC, RV, SGD; Writing: NT, RV, SGD; Resources: CR, FD, MR, PE, RL, RL;

Funding acquisition: DK, SGD.

## Supplementary figure legends

**Figure S1. Identification of Tfh and Tfr and their marker expression patterns.**

(A) Flow cytometry gating strategy to identify follicular T cell (Tfol) subsets (Tfh and Tfr), as well as non-follicular T cell subsets (Tconv and Treg). (B) Illustrations of flow cytometry staining within Tfh from mLNs (top), spleen (middle) and tonsils (bottom) for markers CXCR5, PD-1, ICOS, CD25, HLA-DR, CTLA-4, Ki-67 and CD38. (C) Density plots showing marker expression as logicle-transformed MFI per sample in Tfh (top) and Tfr (bottom) across tonsils (Ton), Spleens (Spl), and mesenteric lymph nodes (mLN) (n=8 each).

**Figure S2. Comparison of IL-1R1 and IL-1R2 expression between follicular and non-follicular T cell subsets within lymphoid organs.**

(A) Example of IL-1R1^+^ flow cytometry staining among Tfh (top) and Tfr (bottom), within mLNs, spleen and tonsils. (B) Illustrations of IL-1R2^+^ flow cytometry staining among Tfh (top) and Tfr (bottom), within mLNs, spleen and tonsils. (C) Percentage of IL-1R1^+^ (top) and IL-1R2^+^ (bottom) cells among CD4^+^ T cell subsets. (D) Density plots showing IL-1R1 and IL-1R2 logicle-transformed expression per sample in Tfh (top) and Tfr (bottom) across tonsils (Ton), Spleens (Spl), and mesenteric lymph nodes (mLN) (n=8 each). **P < 0.01 by paired Wilcoxon test.

**Figure S3. Identification of Tfh, Tfr, and Treg clusters across human secondary lymphoid organs.**

(A) UMAP representations of CD4^+^ CXCR5^+^ (Tfh and Tfr) cells and CXCR5^-^ Foxp3^+^ cells (Treg), overlayed by the expression of the different cell markers. (B) Analysis strategy used for the annotation of follicular T cell and Treg clusters. (C) Frequency of each cluster among all CD4^+^ CXCR5^+^ and CXCR5^-^ Foxp3^+^ cells, across tonsils (Ton), Spleens (Spl), and mesenteric lymph nodes (mLN) (n=8 each). (D) Logicle-MFI of markers used for clustering across the 5 clusters, among all organs. *P < 0.05; **P < 0.01; ***P < 0.001; ****P < 0.0001 by Mann-Whitney test.

**Figure S4. Functionality of the 5 clusters identified following in vitro tonsillar and splenic lymphoid cell stimulation.**

(A) Illustrations of flow cytometry staining for IL-21, IL-2, IL-10 and IL-4 within Tfol from a tonsil (top) and a spleen (bottom). (B) UMAP representations of CD4^+^ CXCR5^+^ (Tfh and Tfr) cells and CXCR5^-^ Foxp3^+^ cells (Treg), overlayed with the expressions of the different cell markers. .

**Figure S5. Modulation of IL-1Rs and IL-1β signaling during in vitro Tfh and Tfr stimulation.**

(A-B) Percentage of IL-1R1^+^ and IL-1R2^+^ cells among Tfh (A) and Tfr (B) within each condition. (C) In vitro Tfr/Tfh cell ratio within each condition. (D) Representation of the IL-1 pathway (WikiPathways WP4496) with differentially expressed genes (DEGs) from scRNA-seq highlighted in red. *P < 0.05; **P < 0.01; ***P < 0.001; ****P < 0.0001 by Mann-Whitney test.

**Figure S6. Correlations between circulating Tfh, Tfr and increase of IL-1β-related molecules within individuals of the Trans*i*mmunom cohort.**

(A) Scatter plot showing the relationship between the number of cTfr and cTfh in the Transimmunom cohort, with the Spearman correlation coefficient and the corresponding p-value. (log2-transformed). (B) MFI measurements of peripheral cytokine and IL-1 related molecules according to the number of autoantibodies in individuals. *P < 0.05; **P < 0.01; ***P < 0.001; ****P < 0.0001 by paired Wilcoxon test.

