## Supplemental data for "IL-1β Signalling Modulates T Follicular Helper and Regulatory Cells in Human Lymphoid Tissues"

Figure S1

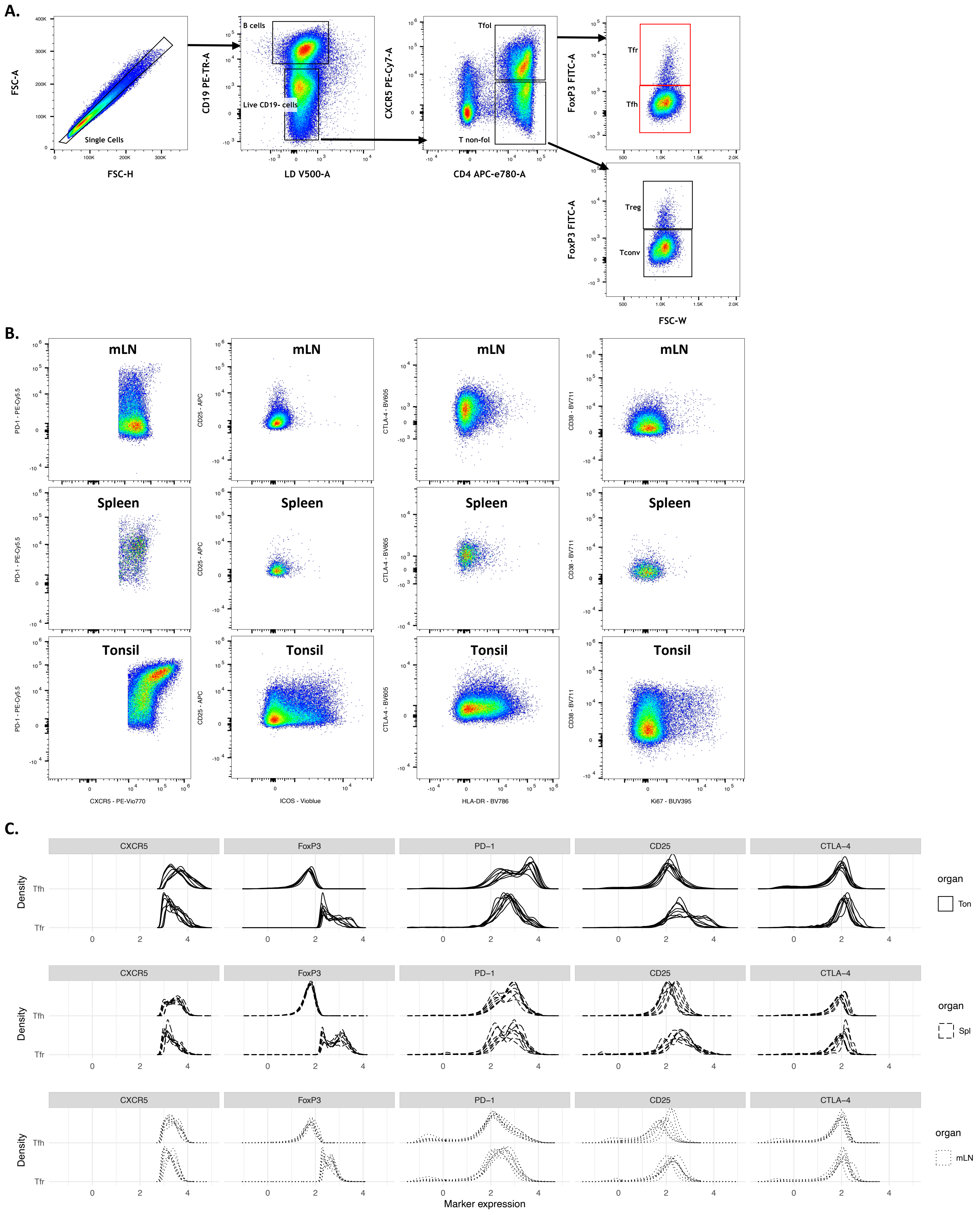
A.
FSC-A
FSC-H
Single Cells
CD19 PE-TR-A
LD V500-A
B cells
Live CD19- cells
CXCR5 PE-Cy7-A
CD4 APC-e780-A
Tfol
T non-fol
FoxP3 FITC-A
Tfr
Tfh
Treg
Tconv
FSC-W
B.
mLN
Spleen
Tonsil
PD-1 - PE-Cy5.5
CD25 - APC
CTLA-4 - BV605
CD38 - BV711
CXCR5 - PE-Vio770
ICOS - Vioblue
HLA-DR - BV786
Ki67 - BUV395
C.
CXCR5
FoxP3
PD−1
CD25
CTLA−4
Density
Tfh
Tfr
organ
Ton
Spl
mLN
Marker expression

Figure S2

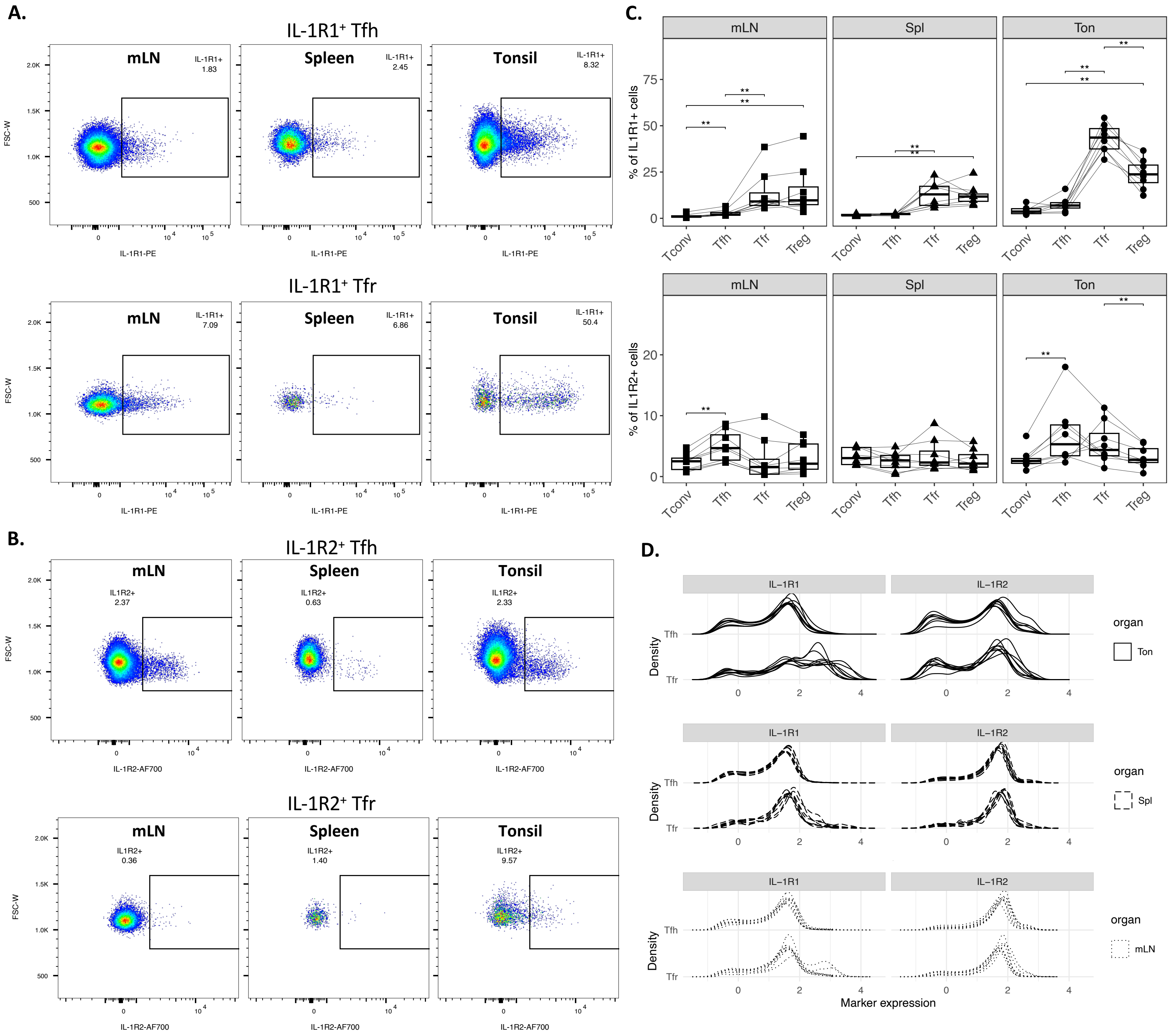
A.
IL-1R1⁺ Tfh
mLN
IL-1R1+ 1.83
Spleen
IL-1R1+ 2.45
Tonsil
IL-1R1+ 8.32
IL-1R1⁺ Tfr
IL-1R1+ 7.09
IL-1R1+ 6.86
IL-1R1+ 50.4
FSC-W
IL-1R1-PE
B.
IL-1R2⁺ Tfh
IL1R2+ 2.37
IL1R2+ 0.63
IL1R2+ 2.33
IL-1R2⁺ Tfr
IL1R2+ 0.36
IL1R2+ 1.40
IL1R2+ 9.57
IL-1R2-AF700
C.
mLN
Spl
Ton
% of IL1R1+ cells
% of IL1R2+ cells
Tconv
Tfh
Tfr
Treg
D.
IL−1R1
IL−1R2
Density
organ
Ton
Spl
mLN
Marker expression

Figure S3

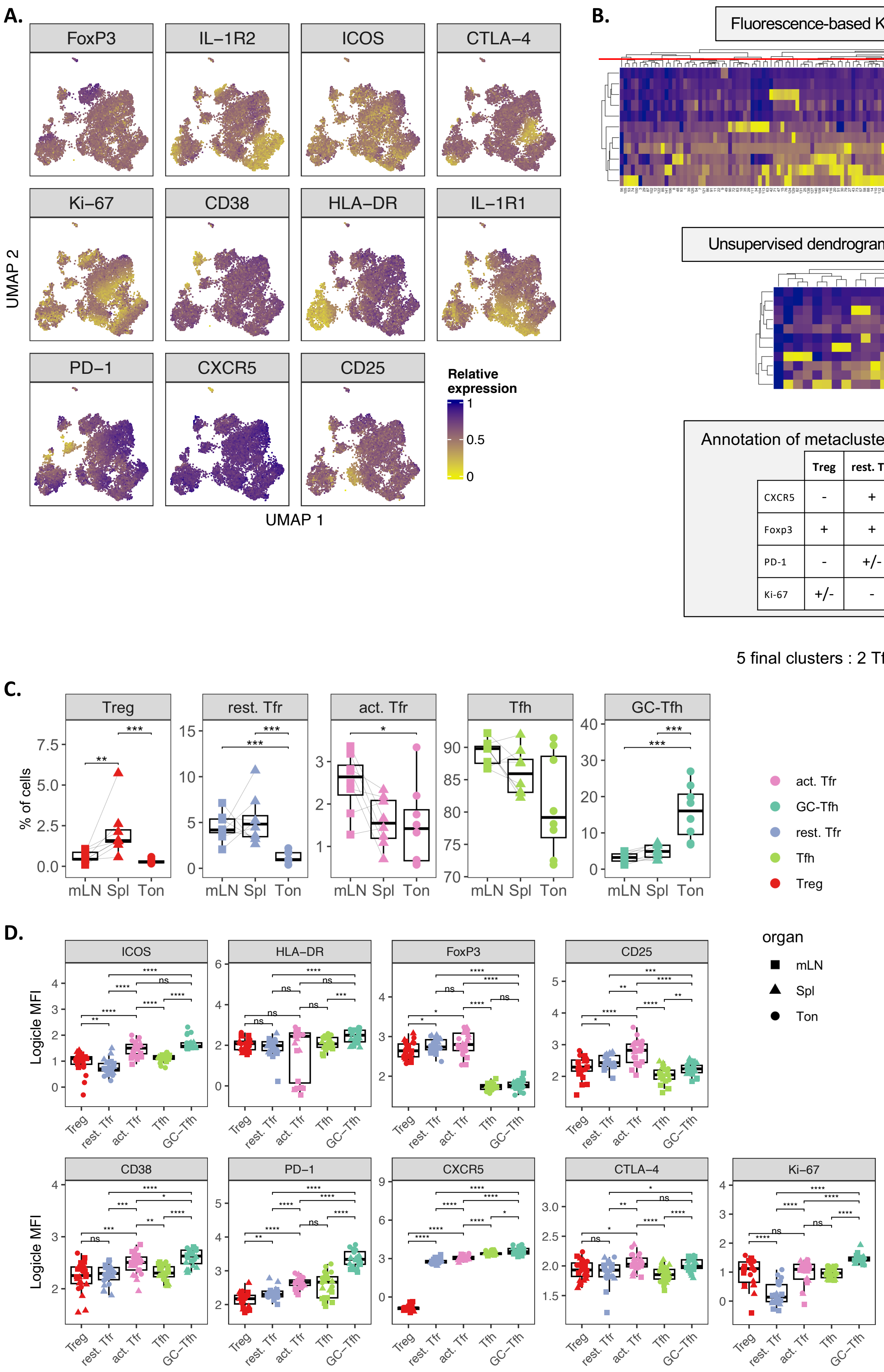
A.
FoxP3
IL−1R2
ICOS
CTLA−4
Ki−67
CD38
HLA−DR
IL−1R1
PD−1
CXCR5
CD25
UMAP 2
UMAP 1
Relative expression
1
0.5
0
B.
Fluorescence-based K-means clustering (n = 150)
Unsupervised dendrogram-based metaclustering (n = 26)
Relative expression
Annotation of metaclusters based on canonical markers
| | Treg | rest. Tfr | act. Tfr | Tfh | GC-Tfh |
| CXCR5 | - | + | ++ | ++ | +++ |
| Foxp3 | + | + | + | - | - |
| PD-1 | - | +/- | + | ++ | +++ |
| Ki-67 | +/- | - | ++ | +/- | +++ |
5 final clusters : 2 Tfh, 2 Tfr and 1 Treg clusters
C.
Treg
rest. Tfr
act. Tfr
Tfh
GC-Tfh
% of cells
mLN Spl Ton
act. Tfr
GC-Tfh
rest. Tfr
Tfh
Treg
D.
ICOS
HLA−DR
FoxP3
CD25
CD38
PD−1
CXCR5
CTLA−4
Ki−67
Logicle MFI
Treg rest. Tfr act. Tfr Tfh GC−Tfh
organ
mLN
Spl
Ton

Figure S4

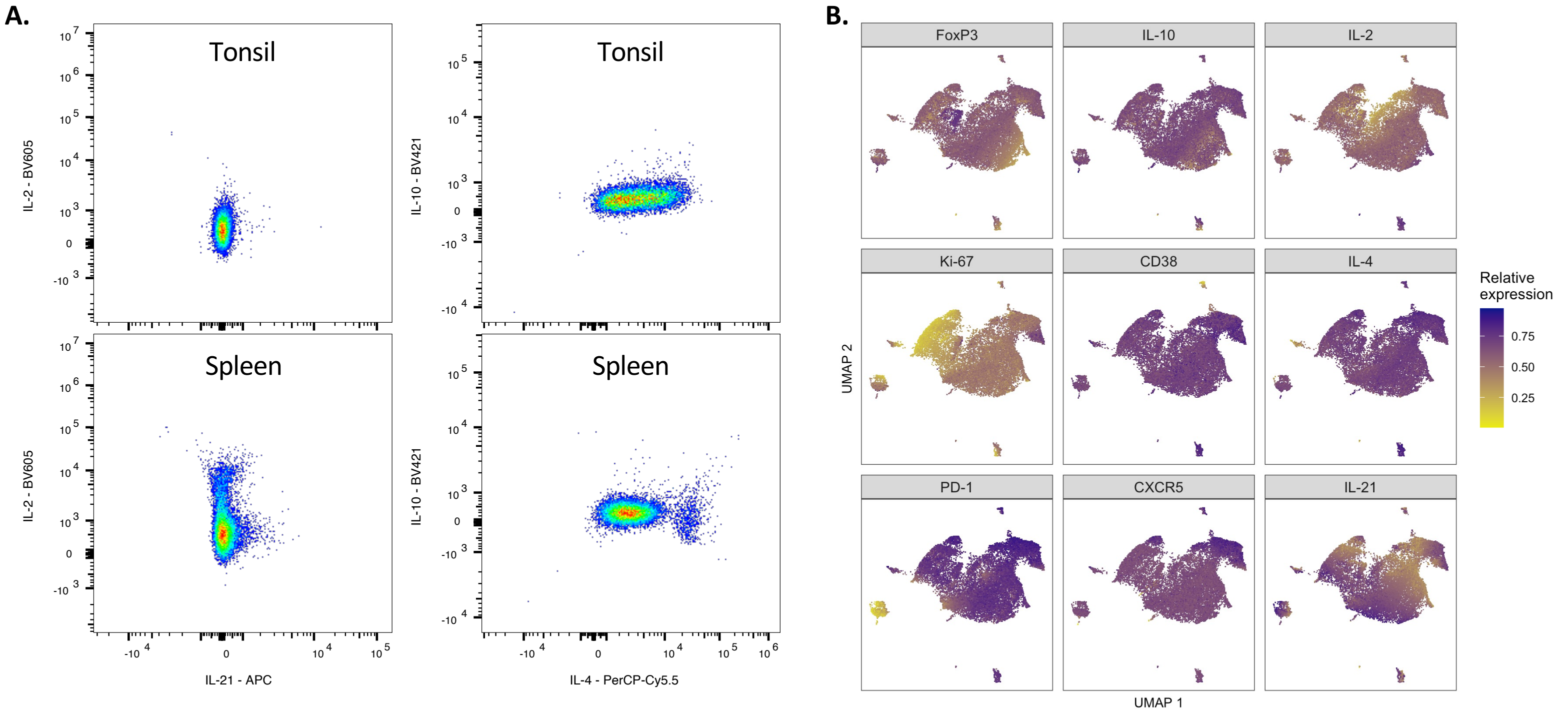
A.
B.
Tonsil
Spleen
IL-2 - BV605
IL-10 - BV421
IL-21 - APC
IL-4 - PerCP-Cy5.5
FoxP3
IL-10
IL-2
Ki-67
CD38
IL-4
PD-1
CXCR5
IL-21
UMAP 1
UMAP 2
Relative expression
0.75
0.50
0.25

Figure S5

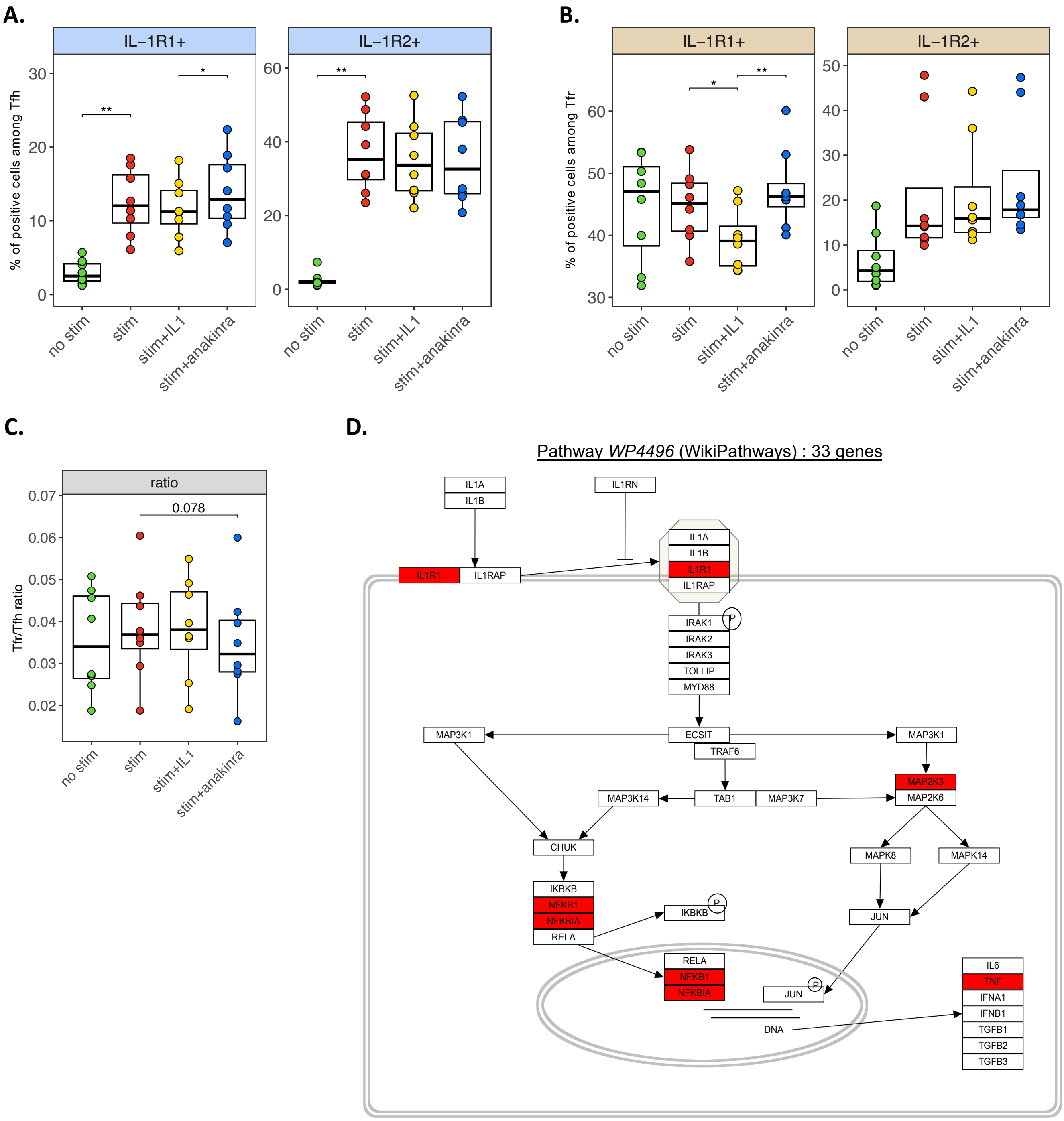


*5 DEG overlapping with 33 genes*

***Hypergeometric test : p = 0.000427***

FIGURE S6

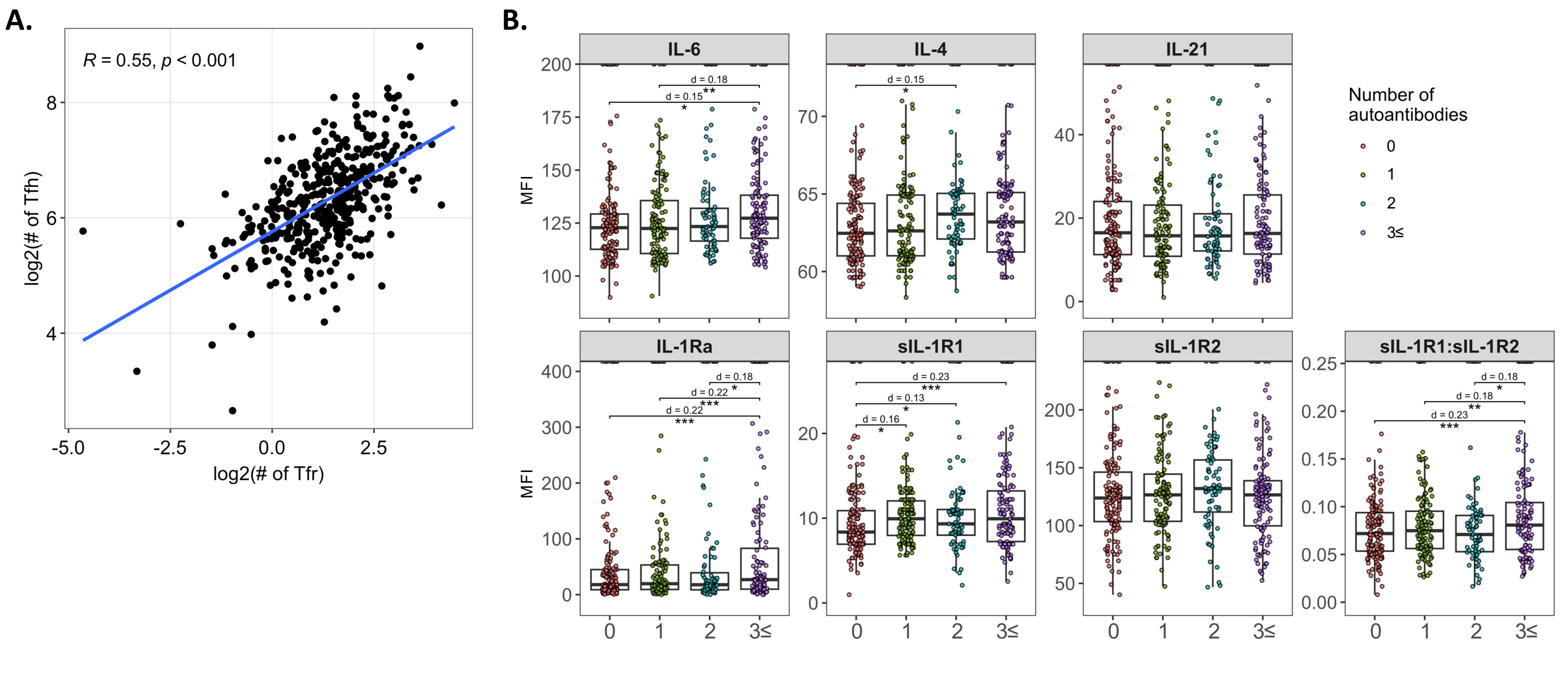
A.
R = 0.55, p < 0.001
log2(# of Tfh)
log2(# of Tfr)
B.
IL-6
IL-4
IL-21
IL-1Ra
sIL-1R1
sIL-1R2
sIL-1R1:sIL-1R2
MFI
Number of autoantibodies
0
1
2
3≤
d = 0.18
d = 0.15
d = 0.22
d = 0.23
d = 0.16
d = 0.13

### Supplementary figure legends

**Figure S1. Identification of Tfh and Tfr and their marker expression patterns.**

(A) Flow cytometry gating strategy to identify follicular T cell (Tfol) subsets (Tfh and Tfr), as well as non-follicular T cell subsets (Tconv and Treg). (B) Illustrations of flow cytometry staining within Tfh from mLNs (top), spleen (middle) and tonsils (bottom) for markers CXCR5, PD-1, ICOS, CD25, HLA-DR, CTLA-4, Ki-67 and CD38. (C) Density plots showing marker expression as logicle-transformed MFI per sample in Tfh (top) and Tfr (bottom) across tonsils (Ton), Spleens (Spl), and mesenteric lymph nodes (mLN) (n=8 each).
